# A Comprehensive Assessment of Methylation-Based Age Prediction Methods

**DOI:** 10.1101/2024.12.23.627443

**Authors:** Han Wang, Jing Qu, Xizeng Zong, Ruirui Cai, Yunkai Li, Shunlin Zhao, Boquan Hai, Ruoting Tian, Zhejun Kuang, Li Zhang, Jian Zhao, Guixia Liu, Chao Zhang

## Abstract

DNA methylation (DNAm) clock is widely used to measure biological age, helping to identify key biomarkers associated with aging, infer the progression of aging, and have promise for elucidating, delaying, or even reversing aging. During the past decade, a large number of epigenetic clocks have been developed. However, they are decentralized, with applicable scopes overlapping. We benchmark 15 of these methods on 142 Illumina DNAm array datasets in five criteria and analyze the biological significance of CPGs about aging and overlapping. There are many exciting commons in models’ performance. We found the optimal model closely related to the numbers and characteristics of the training data. We provided a comprehensive assessment process to guide DNAm clock research at (https://dnamclock.com), the corresponding data and evaluation pipeline are freely available (https://github.comyNENUBioCompute/MethylationEvaluation), this study will aid in the development of improved tools designed to analyze increasingly large DNAm datasets.

## 1 Introduction

Aging is a dynamic and complex biological process reflected in the change of various cellular and molecular processes that accumulate over the course of life, manifesting as impaired functionality and an increased susceptibility to multiple chronic diseases and mortality[1–3]. Several hallmarks[4] are generally recognized as determinants of the aging phenotype and are considered to contribute to elucidating aging. Epigenetic studies are among the most exciting of them all, especially DNA methylation (DNAm), which is one of the most extensively studied and best-characterized epigenetic modifications during aging[5–8].

DNAm is a form of chemical modification in DNA that, unlike sequence mutations, can control gene expression without changing the DNA sequence[2, 9]. The DNAm paradigm plays a crucial role in medicine[10, 11], as specific methylation changes closely correlate with age[12], and these same sites could also play a part in various age-related diseases and cancer[13–15]. Functional studies in model organisms and mammalians suggest that the alterations in DNAm could be utilized to deduce the progression of aging, to identify critical biomarkers related to aging, and to delay or even reverse aging [16–19].

It has been hypothesized that epigenetic age captures some aspect of biological age (BA) and the resulting susceptibility to disease and multiple health outcomes[20]. Hannum et al.[21] and Horvath[22] innovatively utilize the relationship between chronological age (CA) and DNAm changes to measure and compare the aging rate in humans. The approach presents prospects for precise assessments of BA.

During the past decade, extensive effort has been made to develop precise DNAm-based age estimators, also called the epigenetic clock (or DNAm clock). These clocks identify collections of individual methylation sites, with their aggregate methylation status serving as a measure of CA, typically constructed using machine learning (ML) methods[23]. Multifarious DNAm clocks have developed across diverse species, tissue, age spans, and health status [24–35]. However, these clocks are decentralized, with overlapping applicable scopes, and their performance is intricately tied to the constraints of training data, which hampers their convenience and practicality in aging applications.

Although many recent works, such as the Estimage[36] and methylclock[37], attempt to tackle this issue by integrating multiple published DNAm clocks, evaluating and choosing a proper method for a given aging-related investigation is still challenging and problematic. For new field users confronted with an overwhelming choice of methods, it is necessary to make a clear idea of which would optimally solve their problem. For technical developers of the field to make the produced play a valuable role, it is necessary to verify the authority of the existing methods and fully evaluate the advantages and disadvantages. For aging biologists to identify trustworthy biomarkers of aging, it is necessary to verify across multiple factors, such as health status, tissue-specific, life development stage, gender, race, and the datasets used to construct the prediction model. The lack of a comprehensive evaluation and evaluation platform will limit the usage of these DNAm clocks in clinical or biomedical research.

In this work, we focus on human DNAm clocks from birth and compare the performance of the existing and accessible 15 methods, scoring them in 5 criteria while taking into account the phenotypic information of the data. The accuracy, generality, effect of missing value, scalability, and usability of 15 DNAm age estimators were evaluated across 142 Illumina DNA methylation array datasets. Each criterion contains multiple perspectives and is scored by specific metrics (Supplementary Figure 1). We found that no method performed well on all five evaluation criteria, and the optimal model must depend on the numbers and characteristics of the data and be closely related to tissue, age span, and the number of CPG probes. We screened several genes that could potentially serve as aging biomarkers and investigated the impact of disease factors on DNAm clocks. To ensure the broadest reach and maximize the impact of our project, we will release a pre-installed container of the above 15 methods and the performance evaluation pipeline on GitHub to enable future developers to evaluate their methods against others easily. Additionally, we built a web server to allow users to predict their results across multiple methods, which will be convenient for users without sufficient computational skills and resources. We also offer high-quality training-test data for future methodology development, which will be the largest publicly available curated DNAm dataset. The strategy in this work applies to various research fields with extensive studies that can assist researchers in sorting out the research progress and main contradictions in their respective fields and thus can be useful to spearhead the development of new tools.

## 2 Results

### 2.1 Summary

We have selected 15 clocks from existing DNAm clocks methods for evaluation (Fig 1, supplementary Table 1,2). The clocks selected criteria details in the Methods chapter. We collected more than 142 datasets, including 32,595 samples with different tissue types, diseases, age spans, genders, ethnic groups, and platforms (Fig 2.). All samples were manually curated to correct any errors in the data deposited into GEO, including mislabeling, incorrect conversion from raw data, inconsistent annotation with publications, etc. The complete set of curated data is also unified with the same format in array data and metadata, available for free access.

**Fig. 1.**
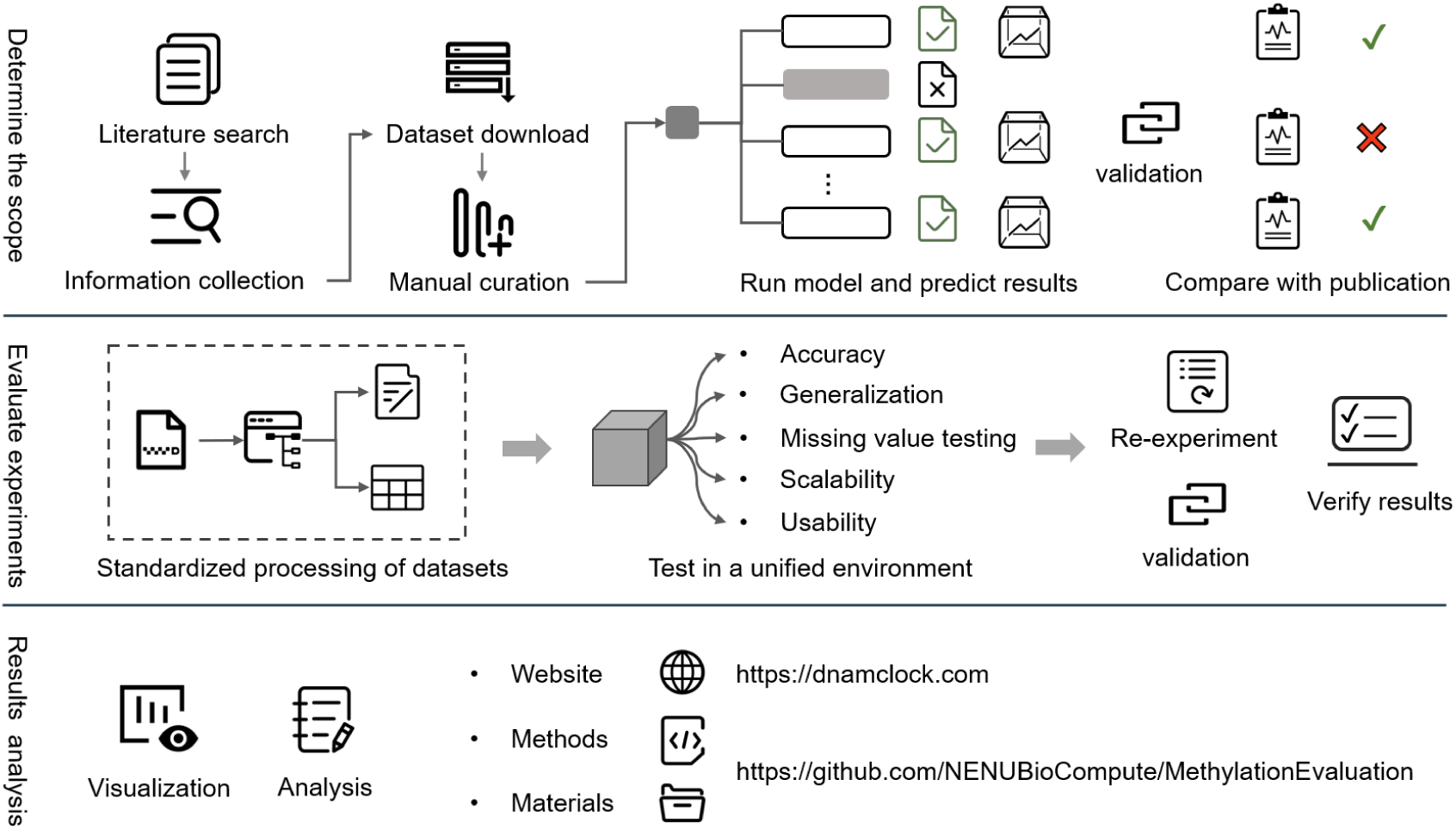
A schematic of overall evaluation pipeline.

**Fig. 2.**
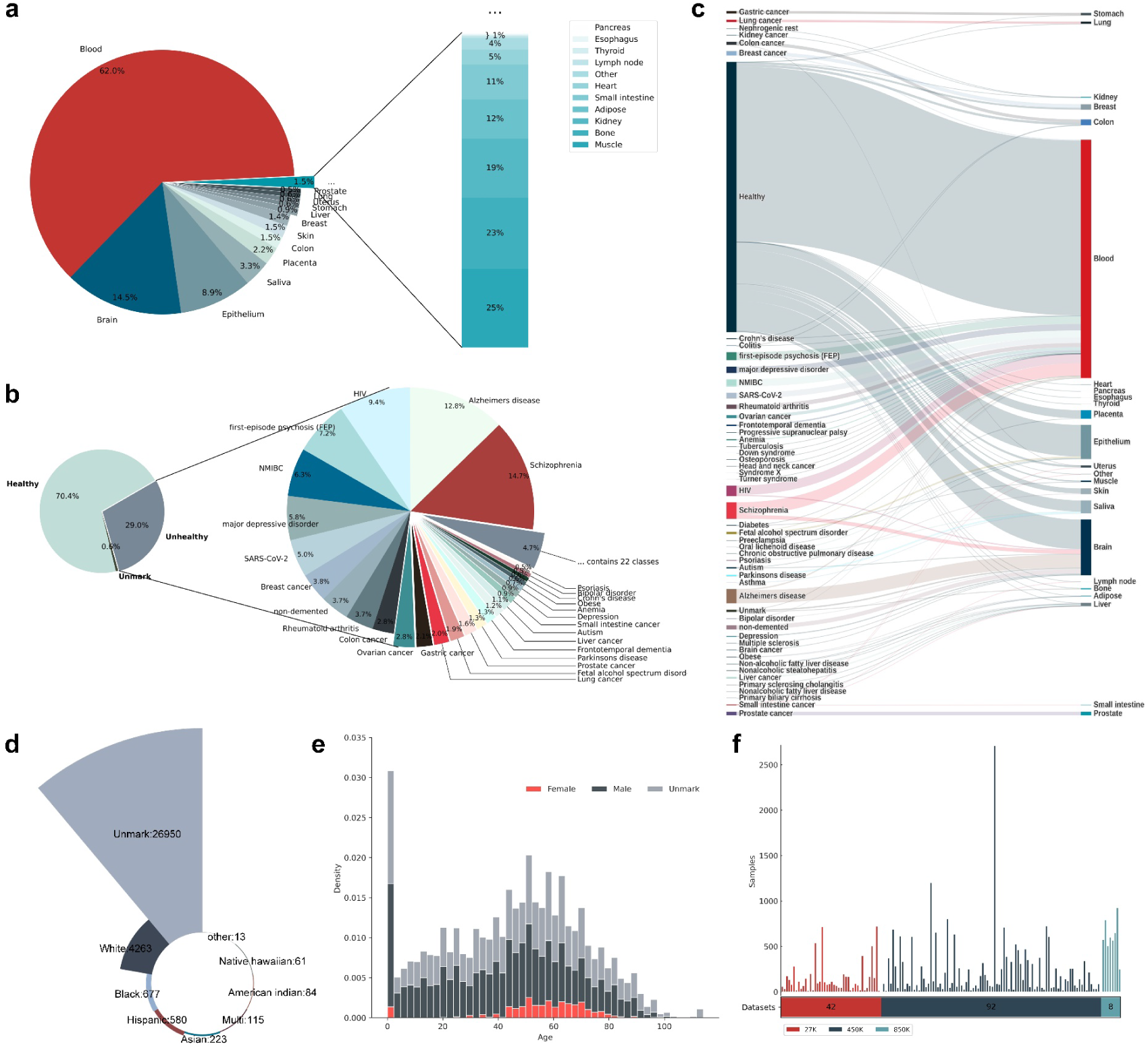
The combination chart presents details from the evaluation datasets. The distribution of a, tissue, and b, disease types. c, The properties relationship of the samples’ tissue and disease. d, The categories and numbers of races. e, The distributions of age and gender. f, the platform to which the dataset belongs.

To compare comprehensively, we established five criteria: (1) accuracy of a prediction on the datasets same as each article, the healthy samples, and the cross-age range samples; (2) generality of the predictions on the cross-tissue samples that were used for modeling or not, and the independent datasets; (3) effect of missing value of model in filled different proportions of CPGs with fixed values; (4) scalability in time and memory concerning the number of samples and CPGs and (5) the usability of the model in terms of code, documentation and the manuscript. Overall, no methods performed well on all five evaluation criteria, but HorvathAge and AltumAge excel in the areas we focus more on (Fig 3.).

**Fig. 3.**
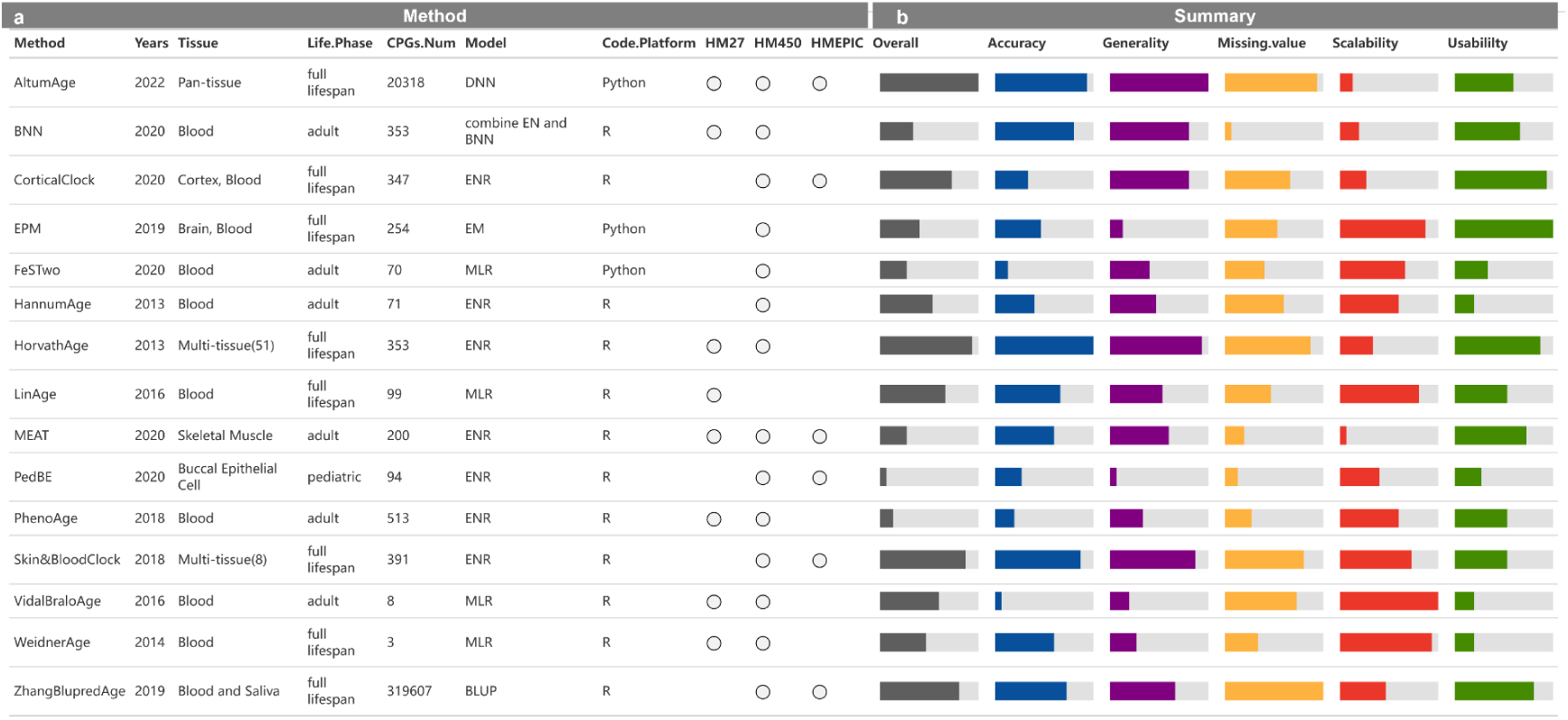
A characterization of 15 methods evaluated in this study and their overall evaluation results. a, Details information of methods. b, Evaluation results span five criteria and the overall ranking combined the five.

More details of each evaluation criterion will be discussed (Supplementary Figure 1), after which we conclude insights for researchers about future perspectives and perform a biological analysis of the CPG sites (CPGs). Finally, we developed website support methods for visual comparison and age prediction.

### 2.2 Accuracy

Accuracy is a metric that measures the degree of proximity between predicted values and actual values, with smaller disparities indicating higher model accuracy, thus reflecting reliability and validity. We evaluated 130 datasets mentioned in each method’s article based on the same data processing.

#### Predictive accuracy in articles

The authenticity and reliability of each method can be illustrated by testing on the samples mentioned in the article (Fig 4.). PhenoAge clock lacks records because its dataset is unavailable. The predicted results are fractionally different from those reported in respective articles, which may be related to the quality or quantity of sub-datasets and batch effect. Overall, a linear relationship between DNAm age and CA can be observed throughout the lifespan. However, significant discrepancies were observed in CorticalClock, EPM, and PedBE. It can be inferred that the predictive accuracy of most methods of testing samples is influenced by the characteristics of the training samples.

**Fig. 4.**
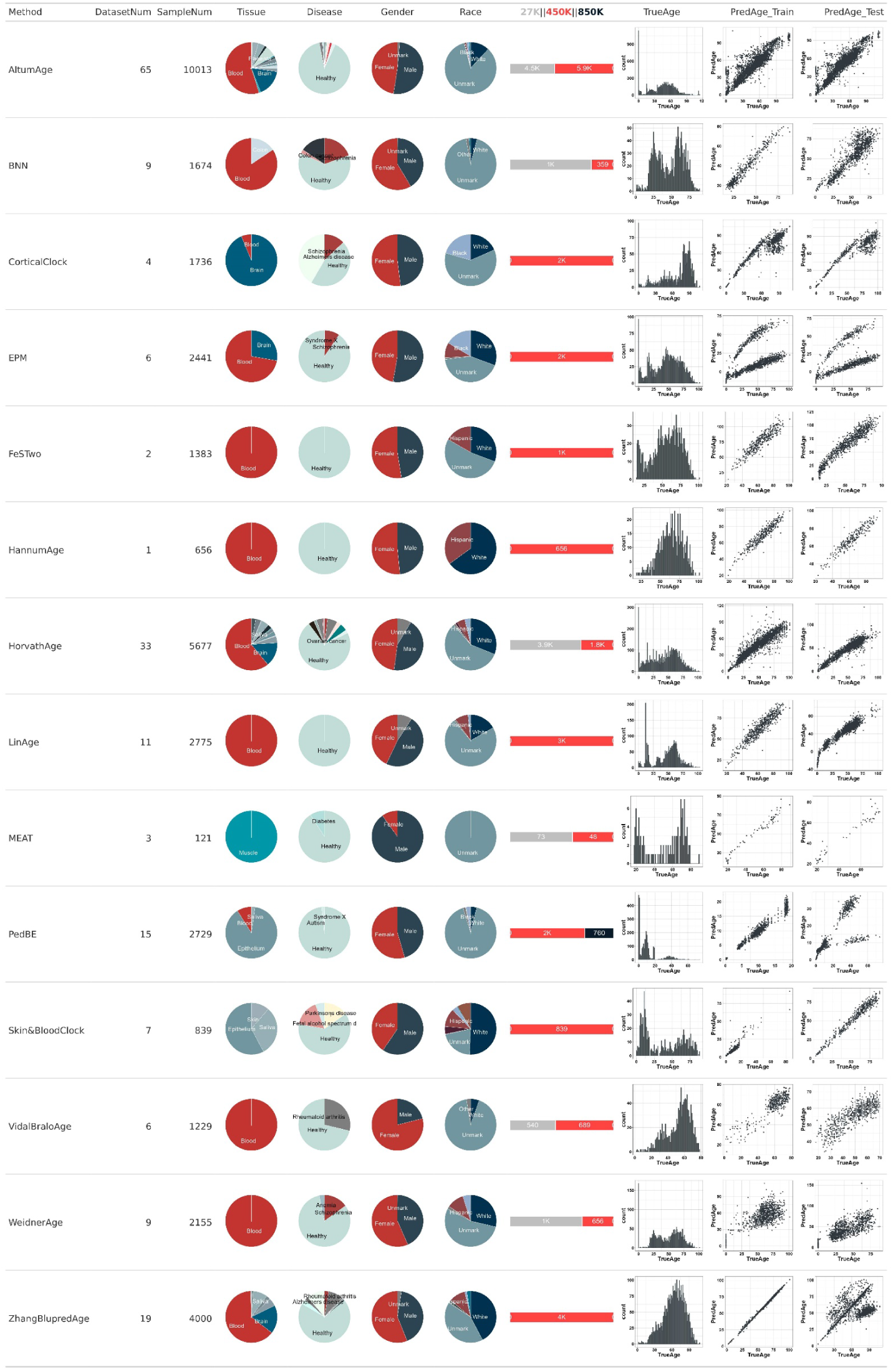
Accuracy in the respective train/test-dataset of model.

#### Predictive accuracy in healthy subjects

Disease may accelerate aging and manifest in increased epigenetic age[36–39]. We depicted the predicted and actual values of 15 methods on healthy samples in 34 datasets that are 450K and contain 51 tissues and age range from 0 to 114 (Fig 5a.). For most methods, a clear trend line was fitted between actual and predicted age. However, a few still produce significant discrepancies. We further calculate the error between each dataset’s predicted and actual values (Figure 5b.). Although there is a combination of factors, the dataset’s age span is key in affecting the predicted accuracy, reflecting each clock’s specific advantage or disadvantage for different age ranges predicted.

**Fig. 5.**
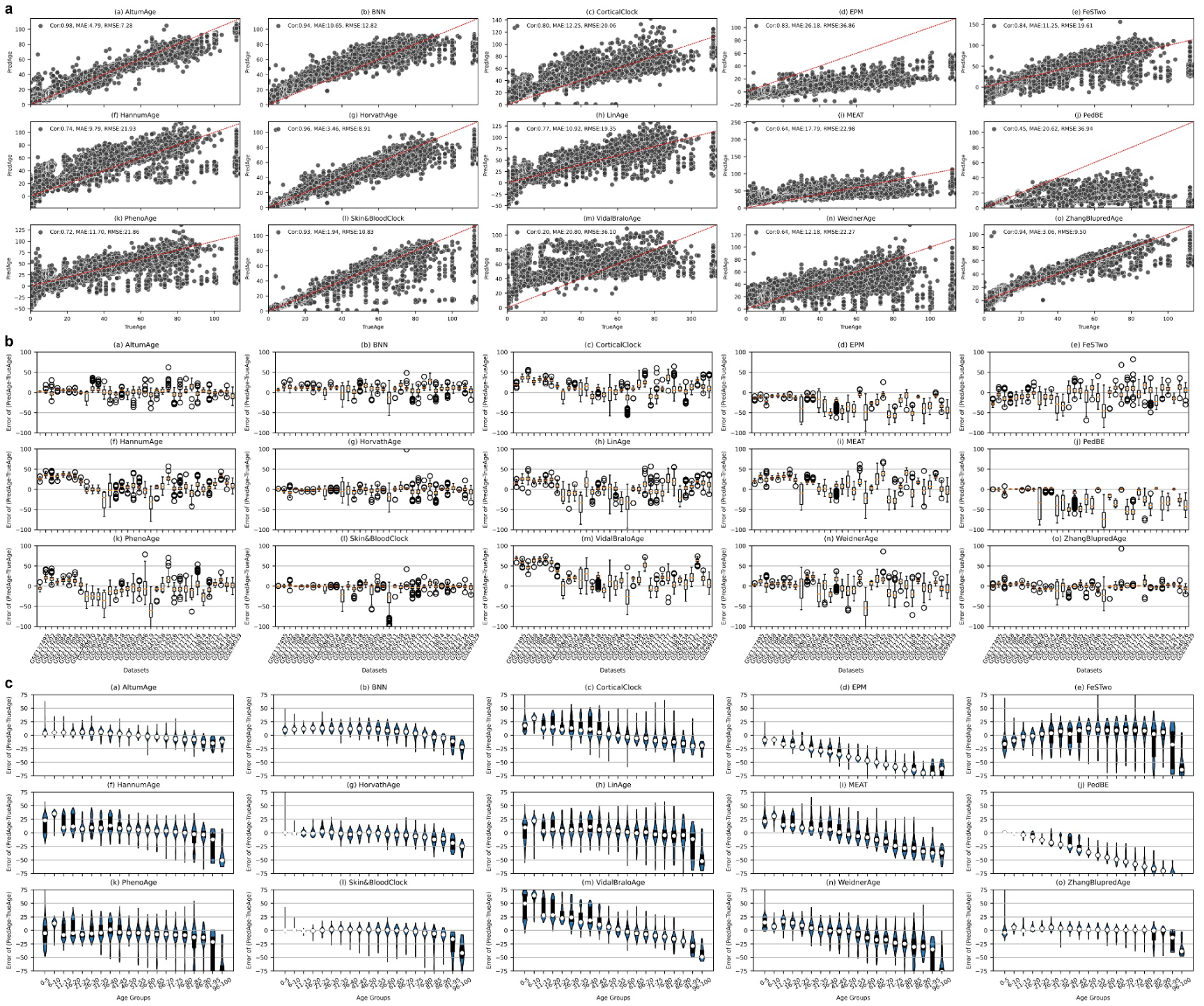
The predicted accuracy of healthy samples, (a)-(o) are shown separately for 15 clocks. a, The distribution trend of predicted and true value in healthy samples of 34 datasets. b, The detailed deviation of DNAm age and CA in each dataset. c, The prediction accuracy of each method at different age ranges.

#### Predictive accuracy across different age-range

It is an established fact that the expression of DNA methylation changes with age[40, 41], as a growing amount of research indicates that the epigenetic clock exhibits different aging patterns between children and adults[42]. We found an obvious diversity in accuracy across different age stages for 15 clocks, with only a few clocks (Fig 5c.), such as PedBE, HorvathAge, and Skin&BloodClock, performing well in younger age groups, and for almost all clocks, prediction accuracy decreased among individuals over 60 years old. It is plausible that the distinct disparities in the aging trajectories across various age spans underlie this phenomenon.

Altogether, the accuracy analysis indicated that existing clocks could reflect the variation trend of DNAm age with CA, but their performance was affected by attributes. However, the predicted accuracy was not stable across datasets, which is significant and complicated and needs to be further resolved. An accurate epigenetic clock for adults over 60 years old is still lacking.

### 2.3 Generality

An epigenetic age estimator is valuable in practice and can be equally compatible with its modeling tissue and other readily accessible human tissue or cells. We executed the generality of each clock on cross-tissue and independent datasets. The average performer of each model was calculated using the same scores to assess the accuracy of prediction (Supplementary Figure 1c).

#### Generality across different tissues

It is the basic requirements of the model that produce an expected result when given input data, similar to training and maintaining stability in unknown data (Fig 6a). The samples we have chosen for testing are both in a healthy state and do not contain any missing CPGs. Details about the age distribution of samples in the three dataset categories for each clock are provided in Supplementary Figure 2.

**Fig. 6.**
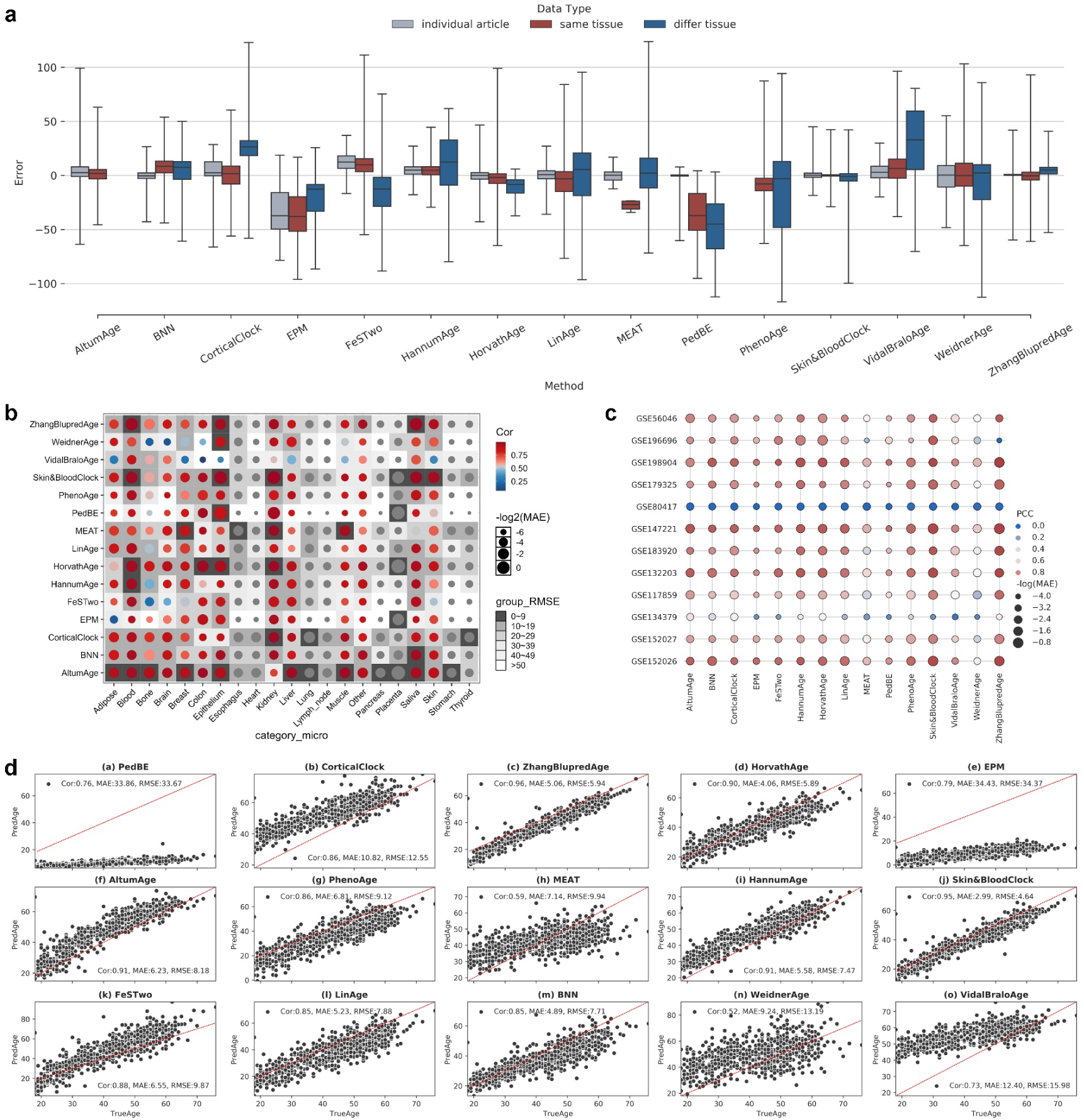
Overview of several key aspects of the generality. a, Generality on three sign data sets: similar with article (red), unsee but similar tissue (blue), and unsee highlight tissue (grey). b, The details of model across tissue. c, the stability and robustness of models in independent datasets. d, The trend of predicted and real value in independent case (GSE132203). The red dashed line at a 45-degree angle represents the ideal trend between predicted and actual values.

The outliers are observed in the predicted outcomes of each clock, which significantly influence the model’s performance. By comparison, HorvathAge, BNN, WeidnerAge, Skin&BloodClock, ZhangBlupredAge, and AltumAge have better robustness than others. PedBE is greatly affected by age range. DNAm clocks can be divided into specific tissue and pan tissue by tissue. Most methods tailored for specific tissues demonstrate higher predictive accuracy within their target tissues than others (Fig 6b). However, some methods exhibit notable performance in non-modeled tissues as well. Pan-tissue clocks that model based on multiple tissue samples exhibit significantly superior generalization to clocks that model specific tissues.

#### Generality on independent data sets

We further predicted and evaluated 12 datasets unknown to all models (Fig 6c). It is noteworthy that the predictive accuracy of the 15 clocks is significantly lower on datasets GSE80417 and GSE134379 compared to other datasets. Details about dataset information can be found in Supplementary Figure 3. Regrettably, a conclusive determination regarding the underlying causes of this phenomenon remains elusive. We used an independent GSE132203 dataset to show the predicted results (Fig 6d).

Overall, the generality analysis indicated that the DNAm clocks can predict well on training samples but have a poor performer for unknown data. The metric score was variable across tissues, and the diversity of training data between methods could indicate that no universal method works well on every tissue. Identifying the underlying causes of outliers and devising appropriate solutions constitutes a potential approach to enhance the model’s predictive accuracy.

### 2.4 Effect of Missing Value

The CPGs value in the expression matrix is missing, which is quite common (Fig 7, Supplementary Table 3). If disregarded the missing, only a limited number of methods can successfully run on test datasets. We simulated the circumstance of model runs after interpolation to evaluate the effect of two imputation values and clock performance (Supplementary Figure 1d). The percentage of missing CPGs increased to 95% from 0%. Given the impact of age distribution on the data, we selected three datasets with relatively uniform age distribution and no missing values for testing. We found that filling with 0 or 0.5 does not significantly impact the model’s predictive performance when the proportion of missing sites is below 10%. The observed trend reveals increased prediction error as the proportion of missing values escalates. We further visualized the results of more collected datasets with missing values (Fig 7.). The predicted ages tend to be the same value in the model of Skin&BloodClock, ZhangBlupredAge, HannumAge, PedBE, FeSTwo, EPM, and CorticalClock due to their CPGs are missing over 70% and even over 90% in 27K datasets. In contrast to other clocks, these exhibit a heightened demand for data platform specificity. Therefore, the CpGs were selected not only to consider their inherent biological significance but also to account for the model’s generalization across multi-platform datasets.

**Fig. 7.**
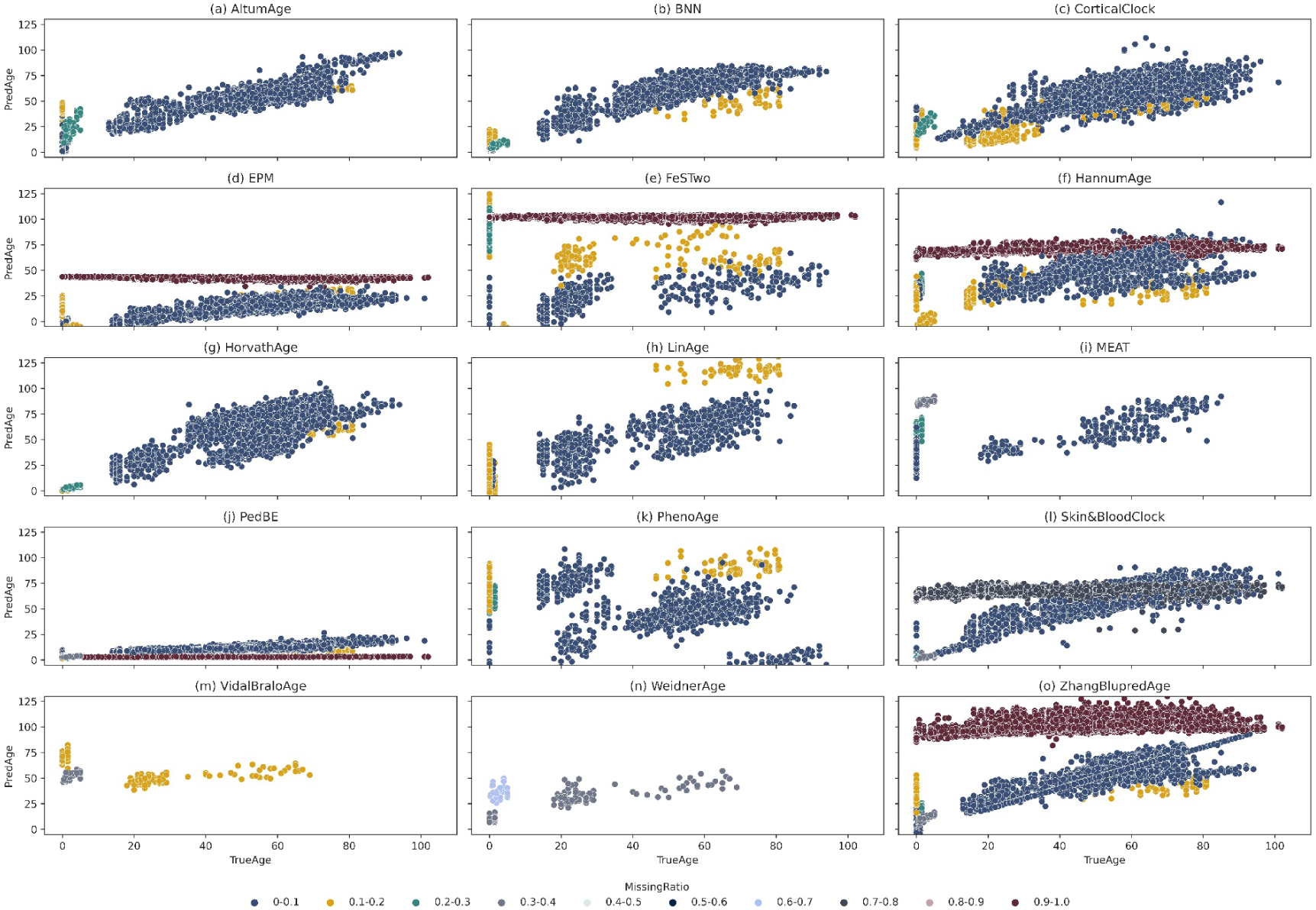
The distribution trend of predicted and real value on datasets with different ratio of missing CPGs. Different color of dot means the degree of missing.

Collectively, the missing value analysis indicated that adjusting the missing value imputation strategy helps improve the model’s predictive performance. As all clocks select a fixed number of CPGs for modeling, the missing proportion can significantly impact the model’s performance and limit its adaptability to cross-platform data. There is no reasonable and effective way to solve when certain CPGs in the dataset are missing in all samples.

### 2.5 Scalability

With the improvement of DNAm sequencing technology, more and more CPG probs are covered. The model should be able to cope with hundreds of thousands of CPGs and as many samples as possible in a reasonable duration. We assessed the scalability of each model from the aspect of running time and memory requirements. Each method was run on up and down-scaled versions of seven distinct real datasets (Supplementary Figure 1e, Time part). The scalability of most methods is generally acceptable.

Data calibration and normalization are the main causes of poor model scalability. Although careful data processing avoids failures during program operation, it also increases runtime and memory usage to some extent. Compared with other models using Elastic Net Regression, the META runs consume more time and memory due to additional data cleaning and calibration. In addition, the more CPG sites the model uses, the greater the cost of processing the sample. Selecting more features for modeling contains more comprehensive evolutionary information but also introduces more interference information and a more significant burden for data processing steps. The running time and memory usage of ZhangBlupredAge based on 319,607 CPG sites increased sharply with sample size (Supplementary Figure 1e, Memory part).

Altogether, the scalability analysis indicated that users should choose suitable models based on the quantity and dimensions of the data, and developers should pay more attention to improving the data calibration and the number of CPGs selected. Effective CPGs can provide potential biomarkers for biologists to cope with aging challenges.

### 2.6 Usability

Although not directly pertinent to the performance of age prediction, it is also imperative to evaluate the implementation’s quality and user-friendliness. We scored each method referring to Saelens’s work[43], including code, documentation, and publication in a peer-reviewed journal.

We found that most methods fulfilled the basic criteria, such as the tutorial and paper quality criteria (Supplementary Figure 1f). However, several methods did not directly provide source code. To reproduce the results, we extensively searched for available code or contacted the authors. Consequently, such methods are not assigned a score in terms of code quality and receive a lower score in terms of code availability.

In general, the usability analysis indicated that the existing DNAm clock is not optimistic about its user-friendliness. Although certain articles detailed the methodology, they need more readily usable code and documents. The design of the DNAm clock in subsequent research needs to be continuously optimized, not only based on the four basic criteria we examine but also more considering users and developers.

### 2.7 WEB Service

Although we have wrapped and made the code public, we still provide users with a website that offers a comprehensive and user-friendly platform for predicting outcomes and analyzing results.

Fifteen methylation clocks were integrated into the website’s online prediction interface, and two functional modules were included.

The first is the evaluation result visualization module, which displays the testing results of the 15 DNAm clocks on 130 standardized DNAm datasets based on the same running environment. Users can view and compare the testing results by selecting datasets and epigenetic clocks. The other is the online prediction module, which supports multiple prediction methods. Users can obtain prediction results by submitting eligible data. Additionally, the website provides a variety of charts, such as scatter plots, box plots, and bar charts, to help users visualize and compare results. The website meets users’ personalized needs and allows them to customize chart styles, colors, labels, and more.

The next step is to integrate more tools, including our creative work. Overall, this website’s characteristics and functionalities will make it a valuable resource for researchers in DNA methylation age prediction.

### 2.8 Biological Analysis for CPGs

To investigate the underlying aging phenomenon and interconnections between DNAm clocks, we analyze the CPGs selected by each model based on overlap and genome. This will provide novel insights into the discovery of aging markers.

The CPGs can be mapped to the corresponding genomic region (Fig 8a). We observed that most CPGs are distributed in the promoter, which may reflect that the genes these CPGs locate change with age. The distribution of CPGs selected by AltumAge’s clock on each chromosome is visualized in Fig 8b. Individual distribution maps for each method are also drawn (Supplemental Figure 4-18.). We observed that few CPGs are selected on sex chromosomes in each method. Several methods suggested that genes on sex chromosomes have different copy numbers and can lead to gender-specific effects on DNAm levels. The distribution of CpG sites on chromosomes can reveal regions rich in methylated sites, which can be used to study changes in DNA methylation levels and gene expression regulation.

**Fig. 8.**
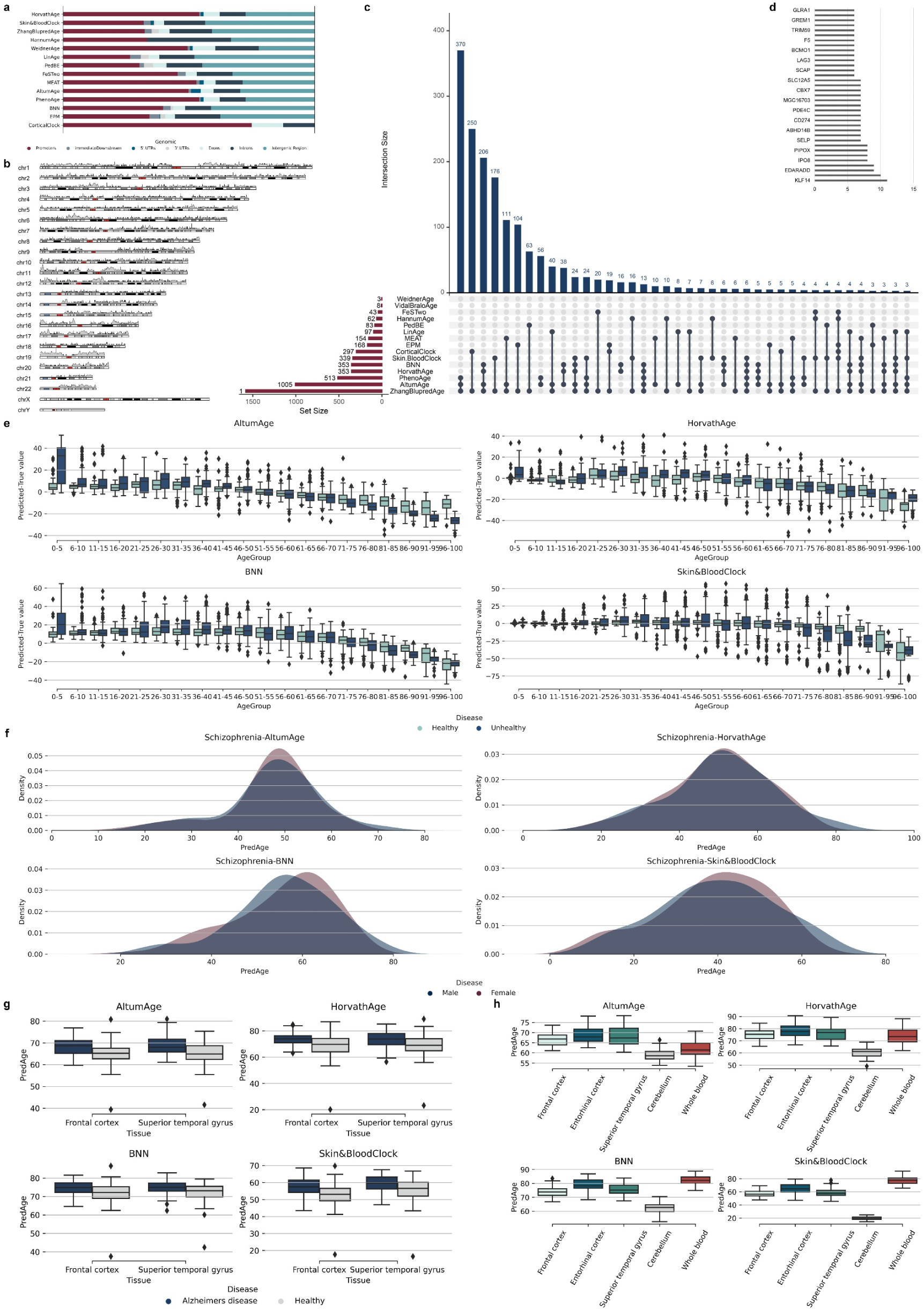
Statistics of several key aspects of DNAm CPG sites. a, The distribution of CPGs over exon, intron, enhancer, proximal promoter, 5 prime UTR and 3 prime UTR. b, Mapping the CpG islands to genome. c, The number of CPGs occurrent in multi-methods. d, Genes were located by ranking the number of co-occurrence CPGs. e, Predicted tends of healthy and unhealthy samples. f, The predicted age distribution of male and female. g, Case of the predicted age difference in the subjects of healthy and Alzheimers disease (using GSE80970). h, Case of the tissue predicted difference in same subject (using GSE59685, AD samples).

Each method’s CPGs have a certain percentage of overlap with each other (Fig 8c). If a gene co-occurs in the selected CpG sites of multiple epigenetic clocks, it may reflect an essential biological function. We mapped the CPGs used in each method to genes and ranked the genes based on the frequency of use across 15 methods (Figure 8d. KLF14 appeared the most frequently, which co-occurrence in 10 epigenetic clocks). KLF14, a master regulator of gene expression in adipose tissue, particularly regulates lipid and glucose metabolism, is associated with the risk of body weight, fat distribution, insulin resistance, type 2 diabetes, and other metabolic syndromes[44]. The NHLRC1 (NHL Repeat Containing 1) and the SCGN (Secretagogin) were found to play a role in the course of certain neurodegenerative diseases[45, 46]. Further, exploring the importance of these genes combined with multi-omics in pathways may guide researchers to breakthroughs in aging.

All in all, the biological analysis for CPGs provides insight for researchers to investigate further the links between genome, aging, and disease. Genes that occur frequently using multiple methods should be further mined and analyzed.

### 2.9 Biological Analysis for Disease

To validate an assumption that the DNAm age of disease samples is older than that of healthy samples, we compared and analyzed the data sets of healthy and diseased samples from the perspectives of tissue, disease, age distribution, and gender. Four reliable models, AltumAge, HorvathAge, BNN, and Skin&BloodClock, demonstrated the age deviation between real and predicted.

The predicted error of the diseased samples was generally higher than that of the healthy samples (Fig 8e). We also observed that most predicted ages are smaller than real ages when individuals are over 60, whether healthy or disease samples. This may indicate that the current clock lacks a comprehensive understanding of age-related changes in the elderly population.

We compared the predicted age of males and females on a data set that was entirely healthy and five types of disease data sets. Most diseases do not reflect significant gender differences (Fig 8f). The variety of predicted age in different tissues of individuals may indicate the different degrees of aging organs or tissue in subjects (Fig 8g, h).

All in all, the difference between the DNAm age of the diseased and healthy sample suggests that the rate of aging and risk of disease in subjects can be inferred based on the DNA methylation clock trained on the healthy sample. However, the existing clocks do not work well for the scheme. In addition, although it is normal for the individual with the disease that DNAm age is older than the chronology age, the reasonable boundary is unknown.

## 3 Discussion

Epigenetics includes modifications to histone proteins, noncoding RNAs, and DNA methylation[47]; here, we focus on the latter since it is a more accessible mark for quantitative measurements. In this work, we presented a comprehensive assessment of 15 epigenetic clocks from 5 criteria based on 142 real datasets and seven synthetic datasets according to corresponding metrics. We focus on comparing the model’s accuracy in healthy samples, the generality of the model in unsee samples, the influence of missing values on the model, the scalability of the model in time consumption and memory usage, and the usability of the model from the user’s perspective.

Based on our benchmark results, we propose a set of practical guidelines for method users and summarize several areas for developers to improve. A convenient website has been developed to support online prediction and evaluation results viewing. Moreover, we summarized genes with high co-occurrence frequency that may be related to aging and diseases. This provides a perspective for researchers to intervene in aging and diagnose and treat diseases. Furthermore, we substantiated the foundational notion that the aging trajectory in healthy samples significantly diverged from that of diseased counterparts. Through a comparative analysis of the disparities in DNAm age between healthy and diseased samples while considering factors such as gender and tissue types.

When estimating an individual’s age using a DNAm clock, several key factors, such as the sample’s health status, age group, and tissue type, need to be considered. The clock selection for prediction can refer to the evaluation report we produce. It is better to check and interpolate missing values in advance to make the prediction more reliable.

Our study indicates that current DNAm clocks are unsuitable for large-scale applications. Their accuracy, generality, and stability in predicting across data sets are limited, and they lack effective solutions for handling missing values and outliers. It is important to balance the impact of data location on time and storage. Furthermore, there is a need for enhanced code sharing and comprehensive functional documentation.

We found that the aging process between tissues is not independent of each other, which is reflected in clocks established based on specific tissues and may also apply to other tissues. While specific models have advantages in certain tissues and age groups, it is increasingly important to develop more versatile models that can effectively differentiate between various tissues and age groups, ultimately delivering better performance.

We summarized several enhancements that could optimize the model’s performance. (1) Deep learning methods exhibit enormous potential for accurately predicting DNAm age. (2) Handling missing values can enhance the model’s generality while employing a reasonable interpolation method can improve its accuracy. (3) The selection of CPGs directly influences the outcomes and interpretability of the model. (4) The resolution of outliers is expected to play a pivotal role, serving as an essential prerequisite for its practical implementation.

In conclusion, future clocks should strategically address the aforementioned issues, aiming to develop superior epigenetic clocks that comprehensively enhance accuracy, generality, stability, scalability, and usability standards. Further, we want to integrate more aging signals into our work, which already has been researched to demonstrate the relationship other for aging[48].

## 4 Methods

### 4.1 Epigenetic clock methods

We searched the literature and gathered a list of DNAm age prediction methods (Supplementary Table 1). Then we excluded the methods from evaluation based on several criteria: 1) not based on DNAm data; 2) does not predict the age of humans from birth; 3) no usable code or website; 4) no programming interface; 5) data of run with cannot available; 6) source code failed to run; 7) does not return an ordering and 8) too slow (requires more than 24 hours on a 100×450K dataset). In the end, 15 methods remained for evaluation: HorvathAge[22], Skin&BloodClock[49], ZhangBlupredAge[50], HannumAge[21], WeidnerAge[51], LinAge[52], PedBE[53], FeSTwo[54], MEAT[55], AltumAge[56], PhenoAge[57], BNN[58], EPM[59, 60], CorticalClock[61], VidalBraloAge[62]. Details can be found in Supplementary Table 2.

### 4.2 Datasets

We collected DNAm data that can be downloaded freely in the public functional genomics data repository Gene Expression Omnibus (GEO). The dataset consisted of Illumina HumanMethylation27 BeadChip (GPL8490, 27K), the Illumina HumanMethylation450 BeadChip (GPL16304, GPL13534, 450K), and Infinium MethylationEPIC (GPL21145, 850K). We included 130 train/test datasets and 12 independent datasets for this evaluation (Supplementary Table 4). Each dataset comprises a gene expression matrix and corresponding phenotypic data. We do standardize for each dataset, respectively.

### 4.3 Filling missing value

We have selected metyhLImp[63] for imputing individual samples missing. However, the method relies on the value distribution of other samples at the same locus for imputation. It is ineffective when all samples in a dataset are missing. A simple imputation approach of filling with fixed values has been chosen in this paper.

### 4.4 Expression Matrix Normalization

Given the difference in the mode of the methylation values in the dataset. There are two formats: (1) the number of methylation and non-methylation or (2) the proportion of methylations to the total. This work adopted the latter form. Therefore, each dataset was checked manually, and the normalization formula (Supplementary Note) was to convert to the latter if it is the former.

### 4.5 Phenotypic Mapping

We standardize the phenotype, including the sample ID, tissue, disease, condition, age, age unit, gender, race, and data platform. The above information was collected from each dataset and underwent the subsequent process: (1) picked field identifier manually for each item from original header file (Supplementary Table 5); (2) extracted and removed the duplicate values under each field based on the identifier; (3) re-defined the name to unify the same means of each field, which is mapping document (Supplementary Table 6); and (4) mapped each phenotypic file to form a new table according to the mapping document.

### 4.6 Data Filtering and Matching

We matched the DNAm expression matrix with the phenotypic data using the sample’s ID. The samples would be filtered out if they had uncharted age, unspecified tissue type, or were unsuccessfully matched.

### 4.7 Benchmark metrics

Age prediction is widely regarded as a regression problem. We selected Pearson correlation coefficient (PCC), median absolute error (MAE), and Root Mean Square Error (RMSE), which are common with age prediction (Supplementary Table 7), to compare the accuracy performance of models. More details about metrics and aggregation strategy are explained in the Supplementary Note.

### 4.8 Method execution

Each method was performed separately in each dataset. The execution is based on the CPU core of an Intel(R) Xeon(R) Platinum 8163 CPU at 2.50GHz, 32G memory capacity, and one R or Python platform was started for each test. During the execution of a method on a dataset, if the memory limit was exceeded or an error was produced, the NA value and error reason were marked for this execution task. Finally, this part is calculated as zero when calculating the total.

### 4.9 Accuracy

We collect available datasets used for model training, testing, or validation in respective articles (Supplementary Table 8), aiming to reproduce results similar to the published ones. Note that the datasets used in our test are not the same as in each article but a subset of them. We set 20 years old as the boundary to divide individuals from birth to the entire lifespan into children (aged 0 to 20 years old) and adults (aged upper 20 years old) (Supplementary Figure 1, Per Age Range: Young, Old.).

### 4.10 Generalization

We divide datasets into training and independent data to evaluate the model’s ability to generality on unknown samples. In order to observe and evaluate the details of cross-tissue, we compare 21 tissues from 130 datasets of articles. Note that all datasets are tested in two cases: unfilled missing sites and filling missing sites (filled with a fixed value of 0.5).

### 4.11 Missing Value Testing

To simulate the missing situation of the CPG site in the dataset, we perform the following step: 1) randomly select a certain percentage of sites from the CPG sites required for the model to run. 2) use the fixed value to cover the selected CPG site for each dataset; 3) feed the imputed data to model as input, and the predictions are output; 4) calculate the error between the CA and predicted DNAm age; 5) repeat steps 2-4 to test the data after filling with 0 and 0.5, respectively; 6) repeat steps 1-5 to test different filling ratios for missing value: 0%, 10%, 20%, 30%, 40%, 50%, 60%, 70%, 80%, 90%, and 95%; 7) repeat steps 1-6 to test each model separately.

### 4.12 Scalability

We created seven datasets and ran only one model at a time on one dataset. A total of 15*7=105 times were performed. The time cost of the model for the run was defined as the duration from the loaded dataset to return predicted results. During code runs, the program is automatically killed when memory usage exceeds the limit, after which we record the error and set it as zero when calculating the score.

### 4.13 Usability

We consider four categories equally important (Supplementary Table 9: Score details table). Each item in each aspect is assigned a weight. The score of each method in each category is calculated by dividing the aggregate score of items by the total score of that category. Finally, the average score of the four categories is used as the model’s final score for usability.

### 4.14 Website

Python 3.8 is the primary development language, while MongoDB is the backend database. Flask is utilized for the backend, and Vue.js is used for the frontend. The interaction between frontend and backend data is achieved through Axios. All online visualizations are implemented using Echarts.js. The project is deployed on the Alibaba Cloud production environment through Nginx and uWSGI to enable online access. Overall, this technical architecture design ensures a robust and reliable platform for users to access a website.

### 4.15 CPGs Analysis

According to the location of the CPG site in the chromosome, we mapped the distribution of all CPGs in each method on the chromosome by the file “450K-hg19” and the R package “karyoploteR”. In addition, we also plot each method separately (Supplementary Figure 4-18.).

We extracted the genes where each method’s CPGs were located from files “450K-hg19” and “450K-hg38”, respectively (Supplementary Table 10). The R package “assignChromosomeRegion” was used to summarize the peak distribution of the genome over exon, intron, enhancer, proximal promoter, 5 prime UTR, and 3 prime UTR. All parameters were used by default. The chromosome range annotation references the ChIP-chip experiments “450K-hg19” and “450K-hg38”.

### 4.16 Disease DNAm Age Analysis

The data for analysis was selected from the 450K platform datasets that cannot miss the CPGs required for the four methods. The healthy data were fetched from the datasets, and all samples were recorded as “Healthy,” which means if the dataset has an unhealthy sample, that will not be in the selection. The disease data were selected from samples with disease records, whether the dataset contained healthy samples or not. More analysis on other diseases and datasets are presented in the supplementary Figure 19-27.

## Funding

This work was supported by the National Natural Science Foundation of China (No: 62372208, 62372099); the Jilin Scientific and Technological Development Program (No. 20230201090GX); and the Fundamental Research Funds for the Central Universities (2412023YQ002).

## Acknowledgments

We would like to thank Hanlin Gao, Wei Yao, Liming Guo, and Simin Liu for validating the datasets referenced in the evaluated articles. We also appreciate Kangwei Geng’s help with developing the website interface.

## Competing interests

The authors report no competing interests.

## Data availability

The list of all processed real and synthetic GEO datasets used in this study is deposited on the GitHub repository (https://github.com/NENUBioCompute/MethylationEvaluation). GitHub also links to a Google Drive where our gathered DNA methylation data is publicly available. A summary of the results reported can be found on the website (https://dnamclocks.com).

## Code availability

The run scripts call several R and Python packages. Detailed instructions on how to use them can be found in the GitHub repository (https://github.com/NENUBioCompute/MethylationEvaluation). The repository deposited all code for this work, from data processing to results analysis.

## Notes

### Competing Interest Statement

The authors have declared no competing interest.

