## Supplementary Figures for "A Comprehensive Assessment of Methylation-Based Age Prediction Methods"

### Supplementary Figure 1

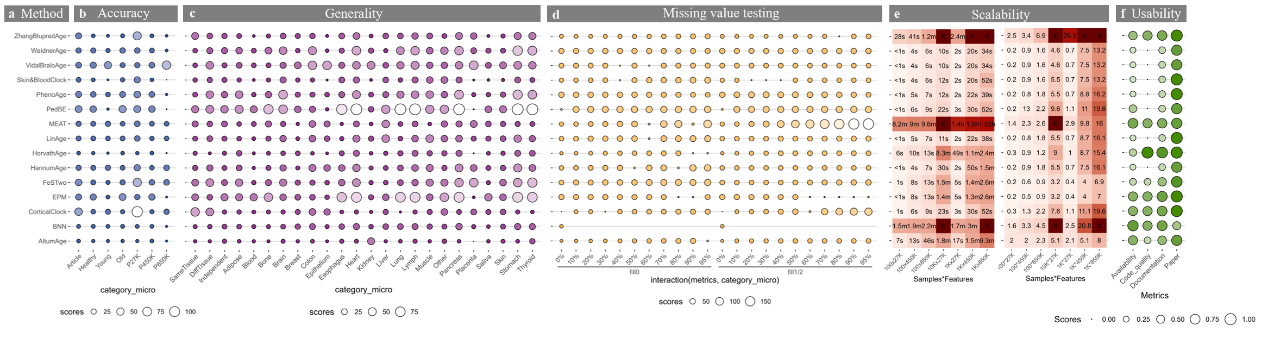

Fig.3 Detailed results of the five main evaluation criteria: accuracy, generality, missing value testing, scalability, and usability. The larger and darker the bubble indicates that the model's prediction error is smaller. We plot the reciprocal of RMSE in accuracy, generality, missing value testing. a. The names of the methods, ordered as in Fig. 2. b. Accuracy of methods on healthy samples across tissue, and age range dataset. We only used datasets from 15 articles to evaluated in this part. c. Generality results by calculating the RMSE score of the model on unknown samples. These samples can be divided into three data sets: from 130 data sets, the unknown samples for each model are divided into datasets of the same and different tissue as when the model was trained/tested; a separate test set that is completely independent of 15 articles. d. The impact of different imputation methods on model performance and the stability of the model under different missing conditions measured by RMSE. The testing involves two imputation methods: 0 and 0.5, each constructed with 10 missing cases. e. Predicted execution times and memory usage for varying numbers of samples and CPG sites (no. of samples × no. of cpgs). Running each method on seven synthesized datasets with real data. k, thousands; m, millions. f. Usability scores of the tool and corresponding manuscript, grouped per category.

### Supplementary Figure 2

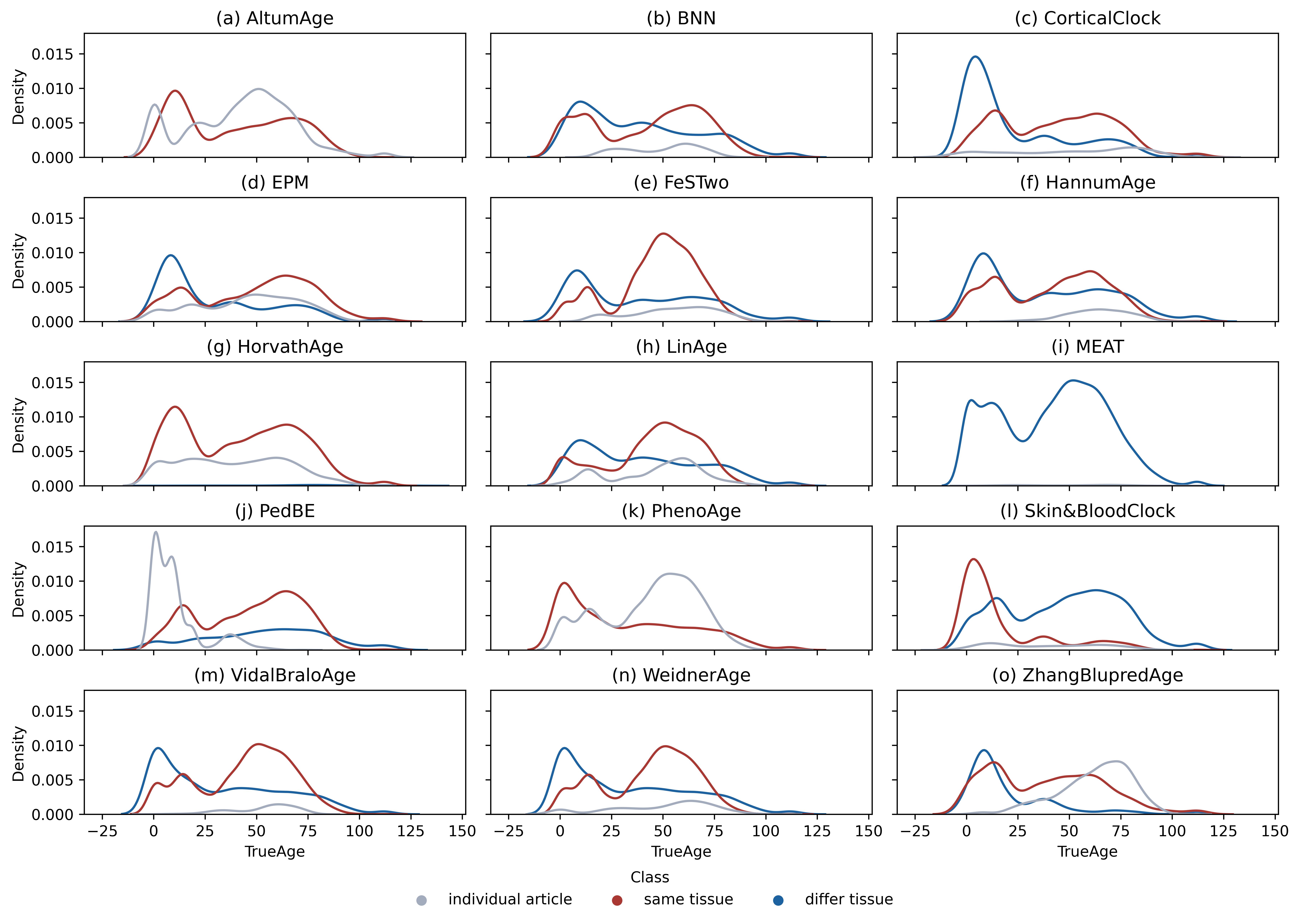

### Supplementary Figure 3

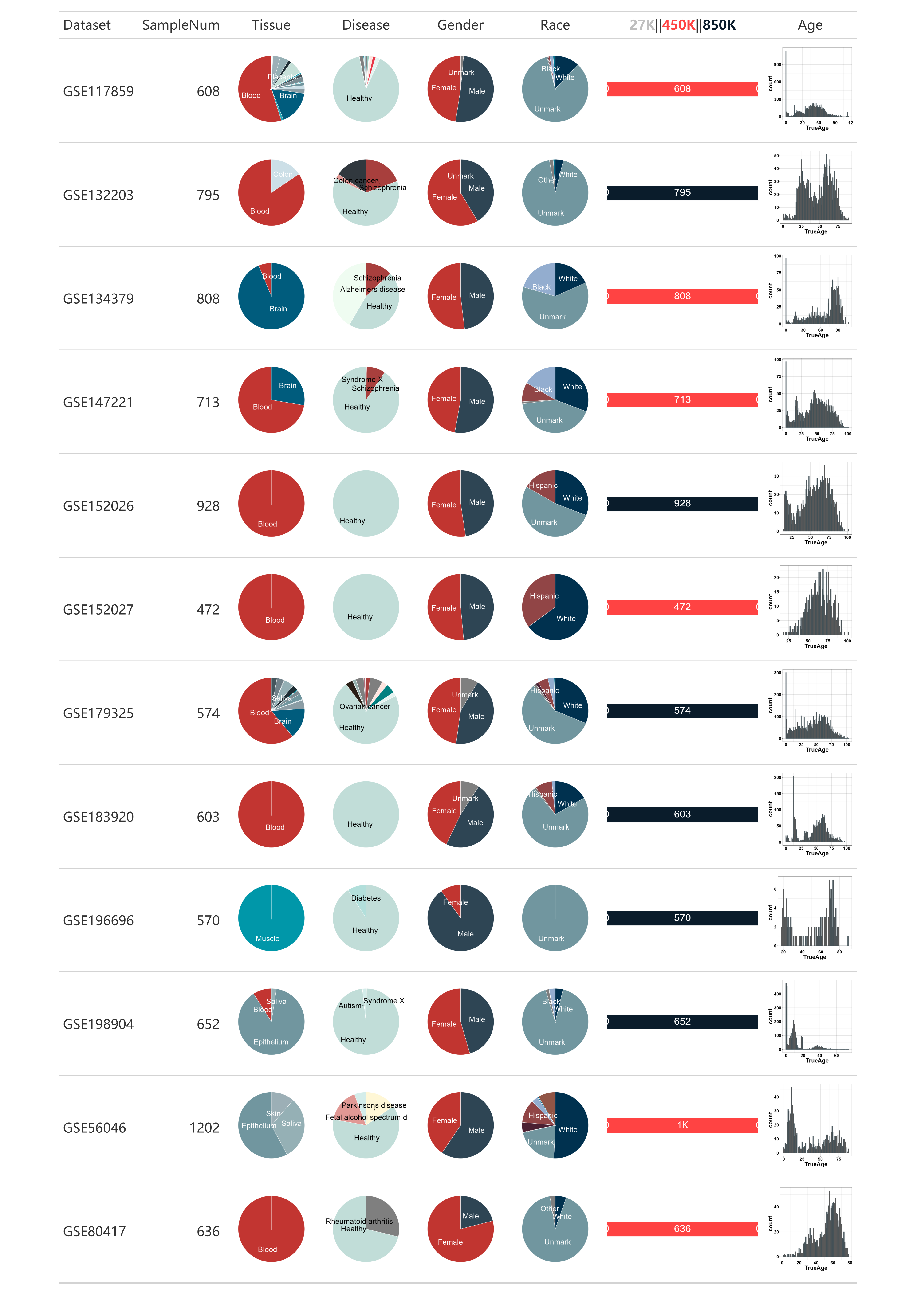

### Supplementary Figure 4: HorvathAge

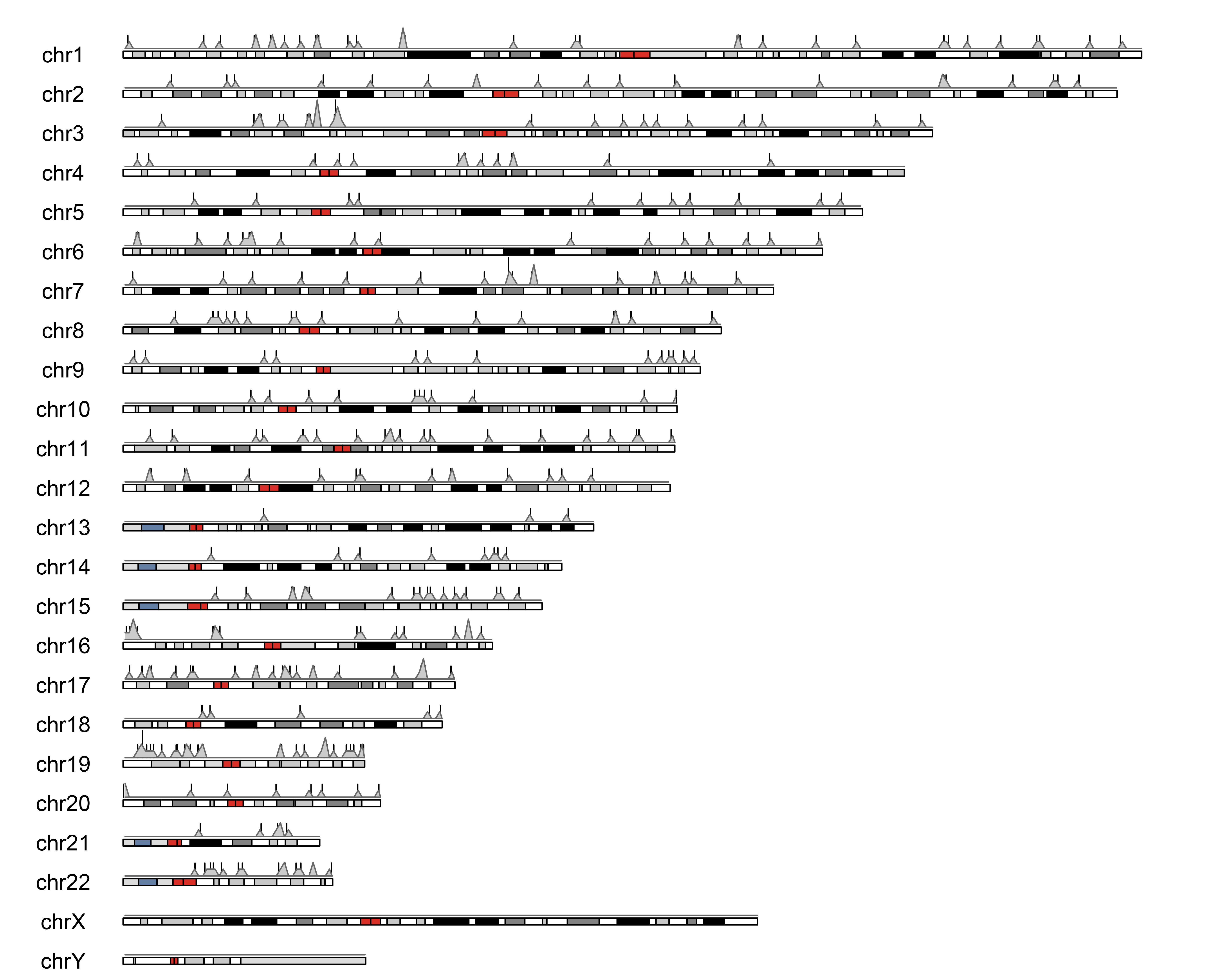

### Supplementary Figure 5: Skin&BloodClock

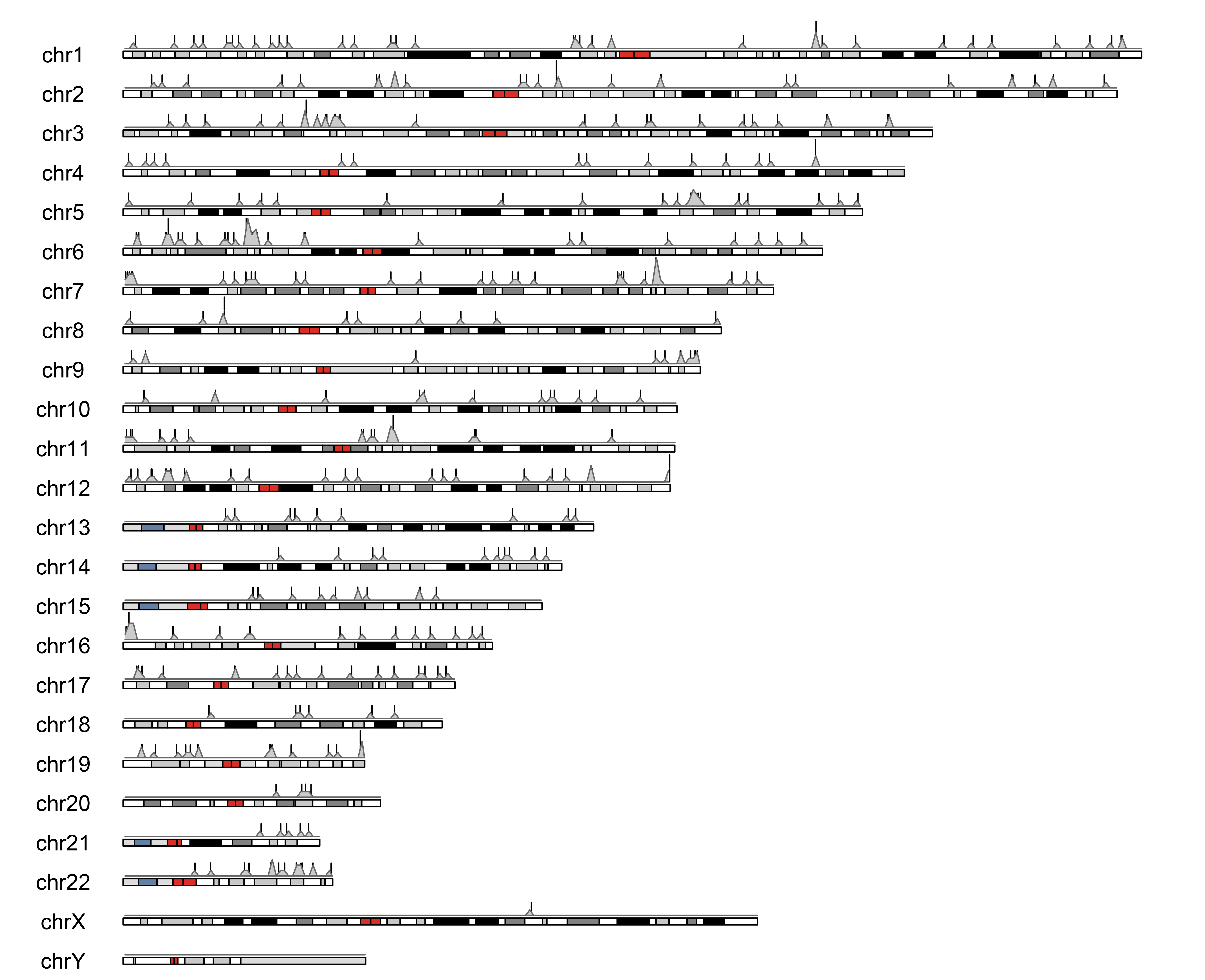

### Supplementary Figure 6: ZhangBlupredAge

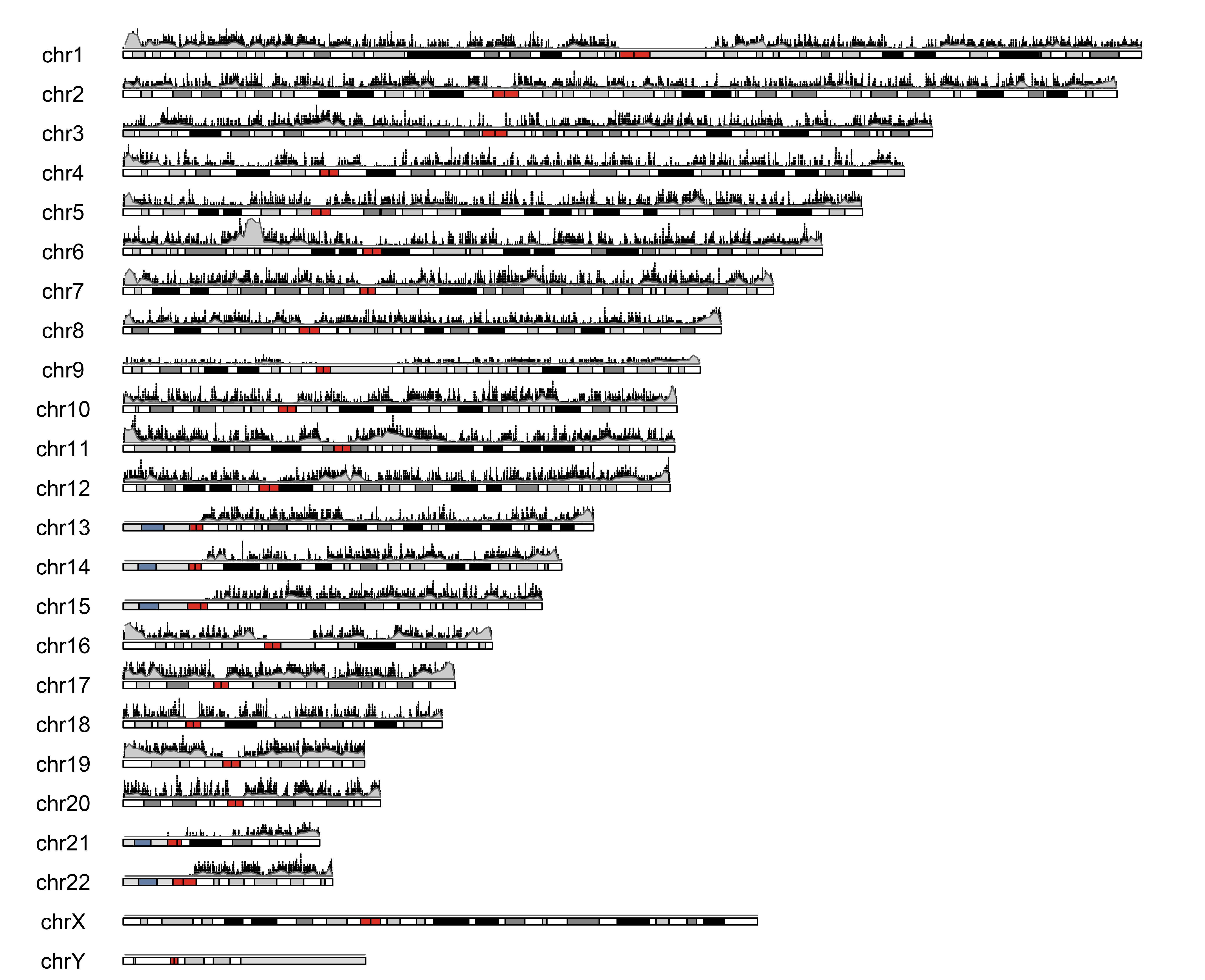

### Supplementary Figure 7: HannumAge

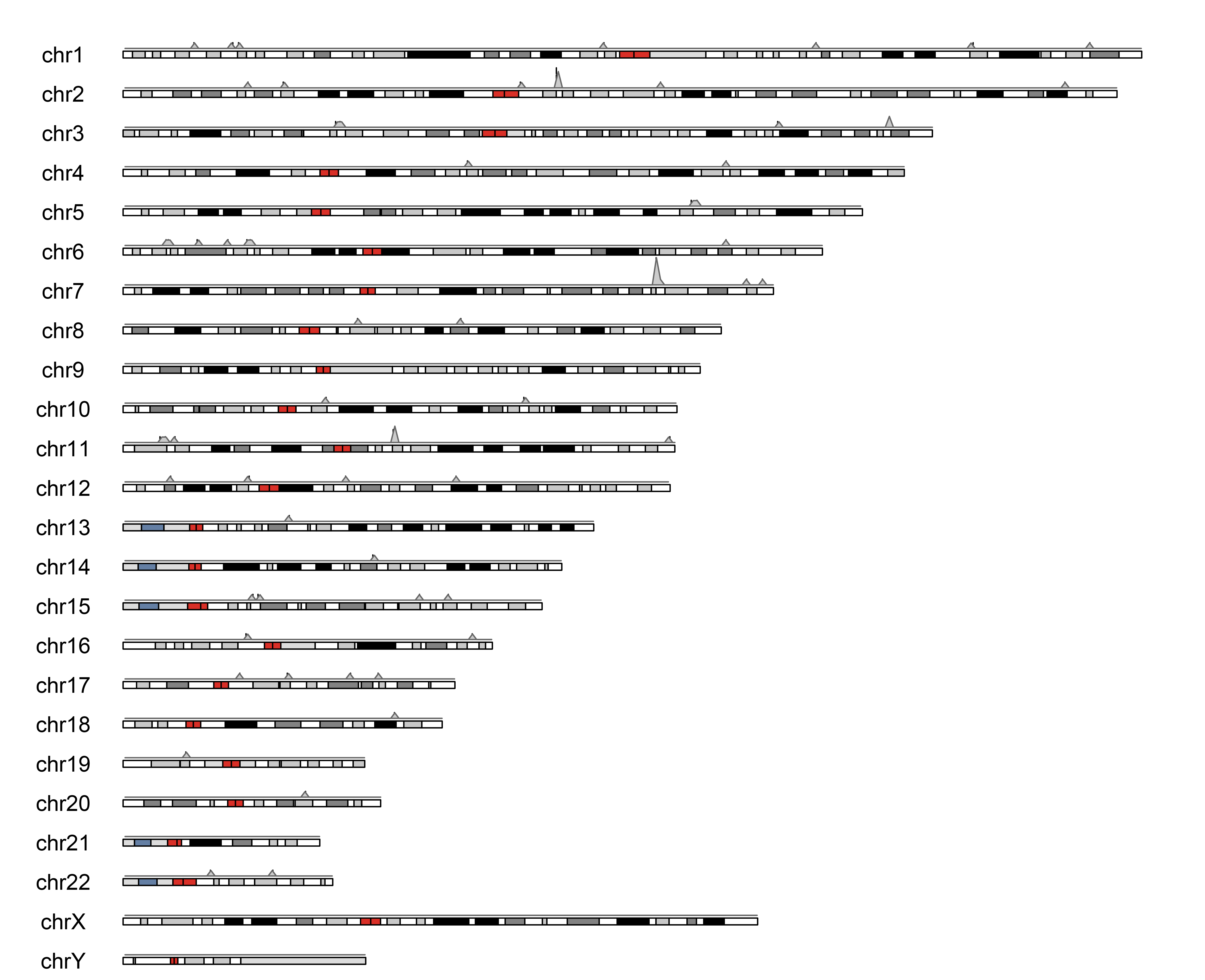

### Supplementary Figure 8: WeidnerAge

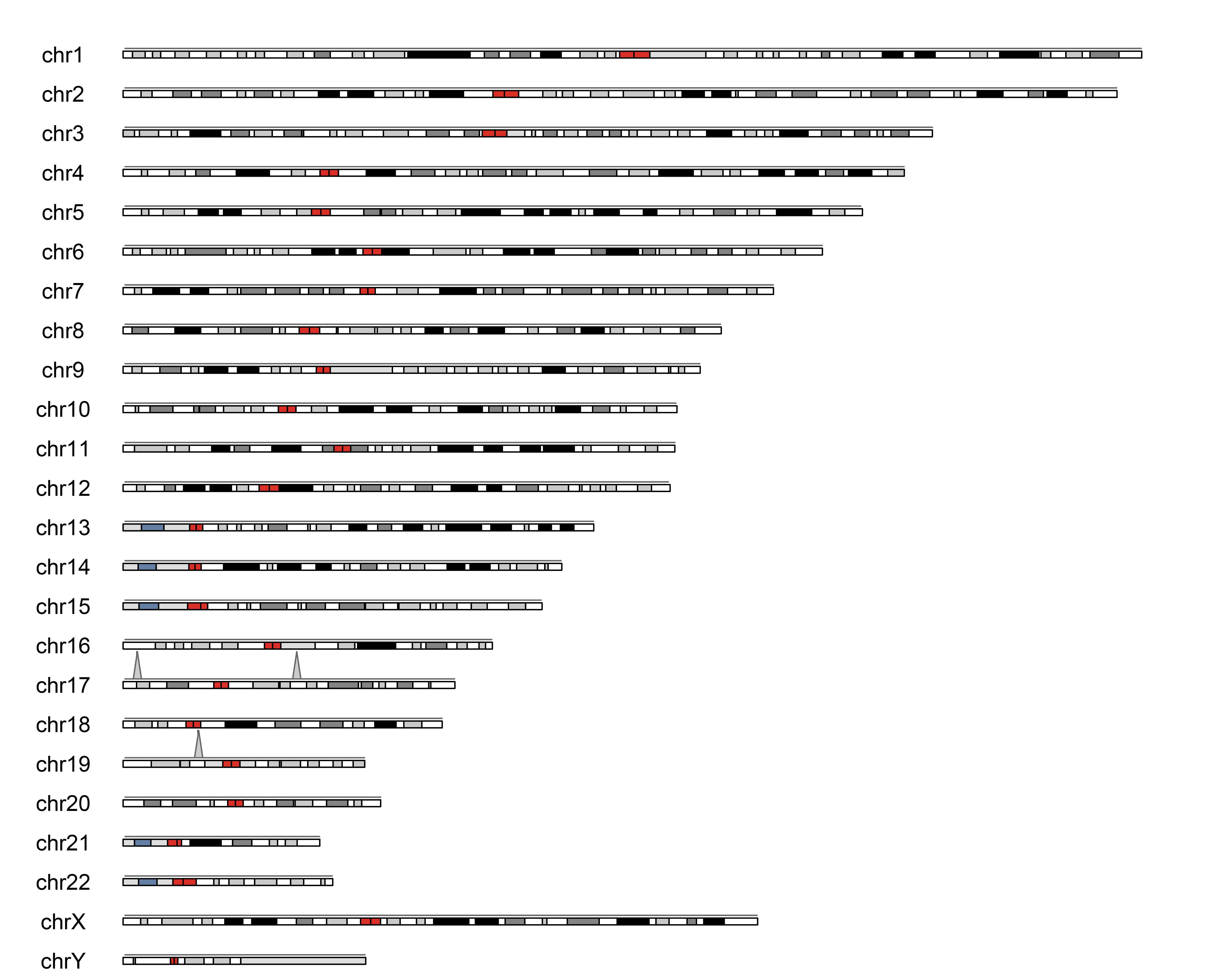

### Supplementary Figure 9: LinAge

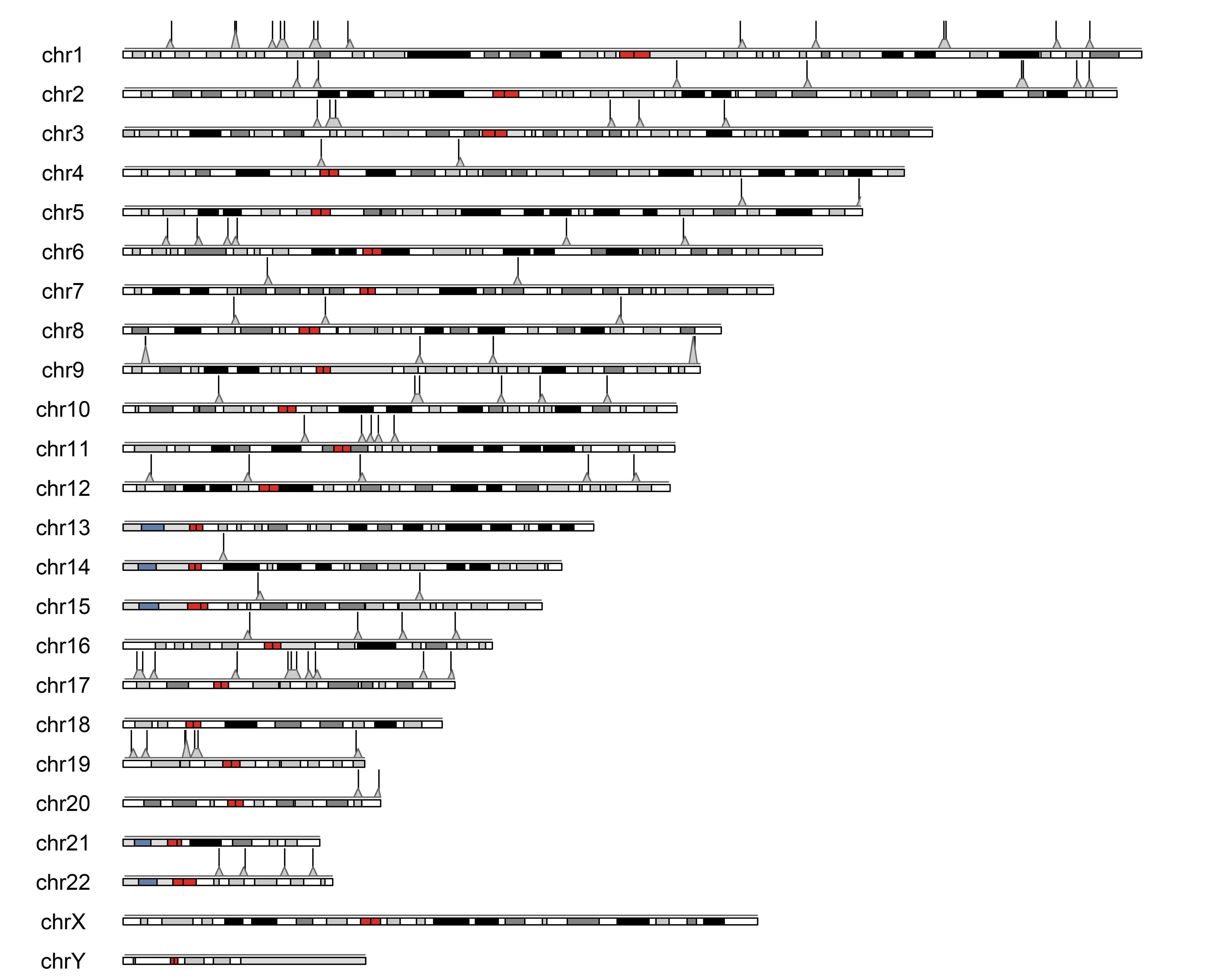

### Supplementary Figure 10: PedBE

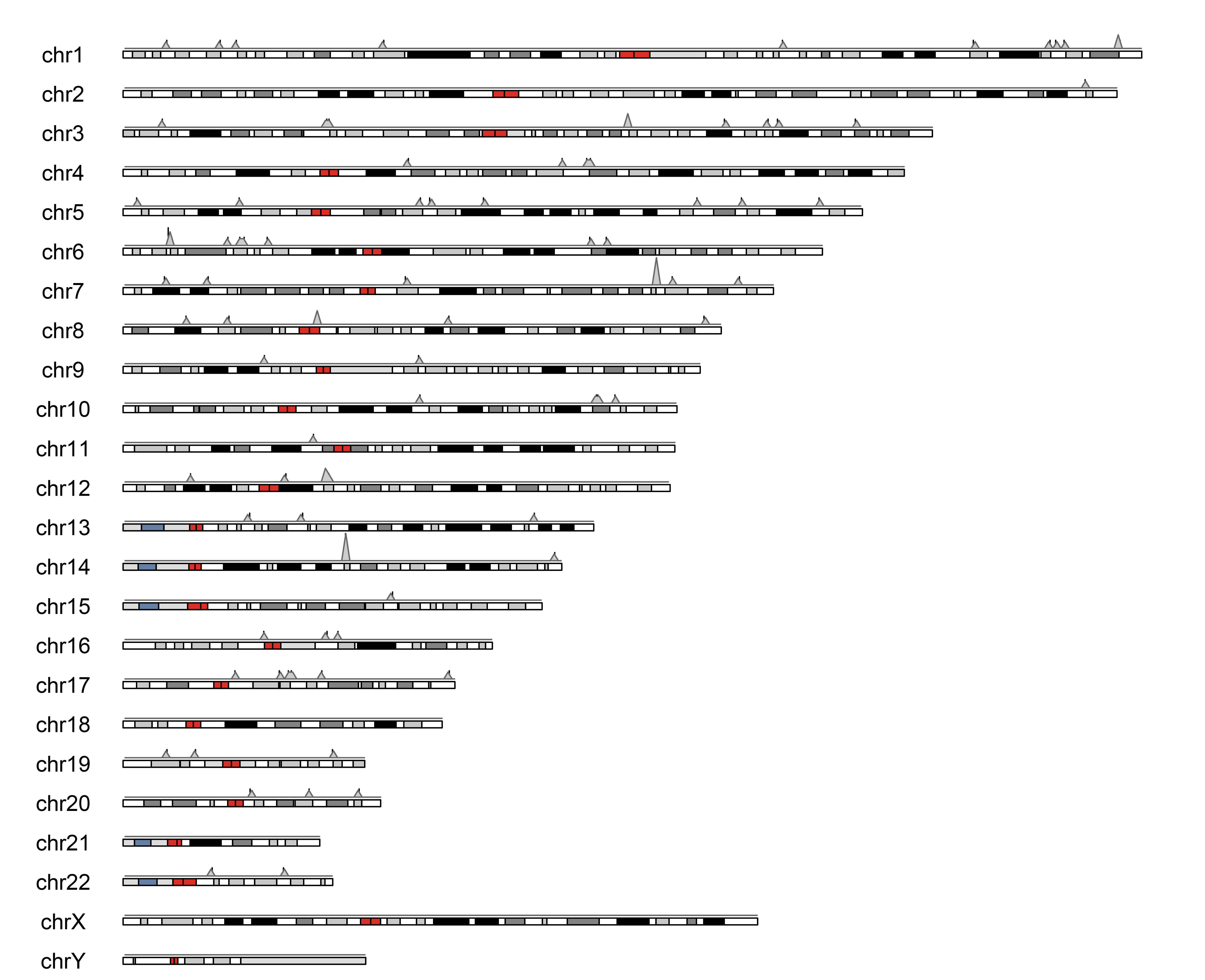

### Supplementary Figure 11: FeSTwo

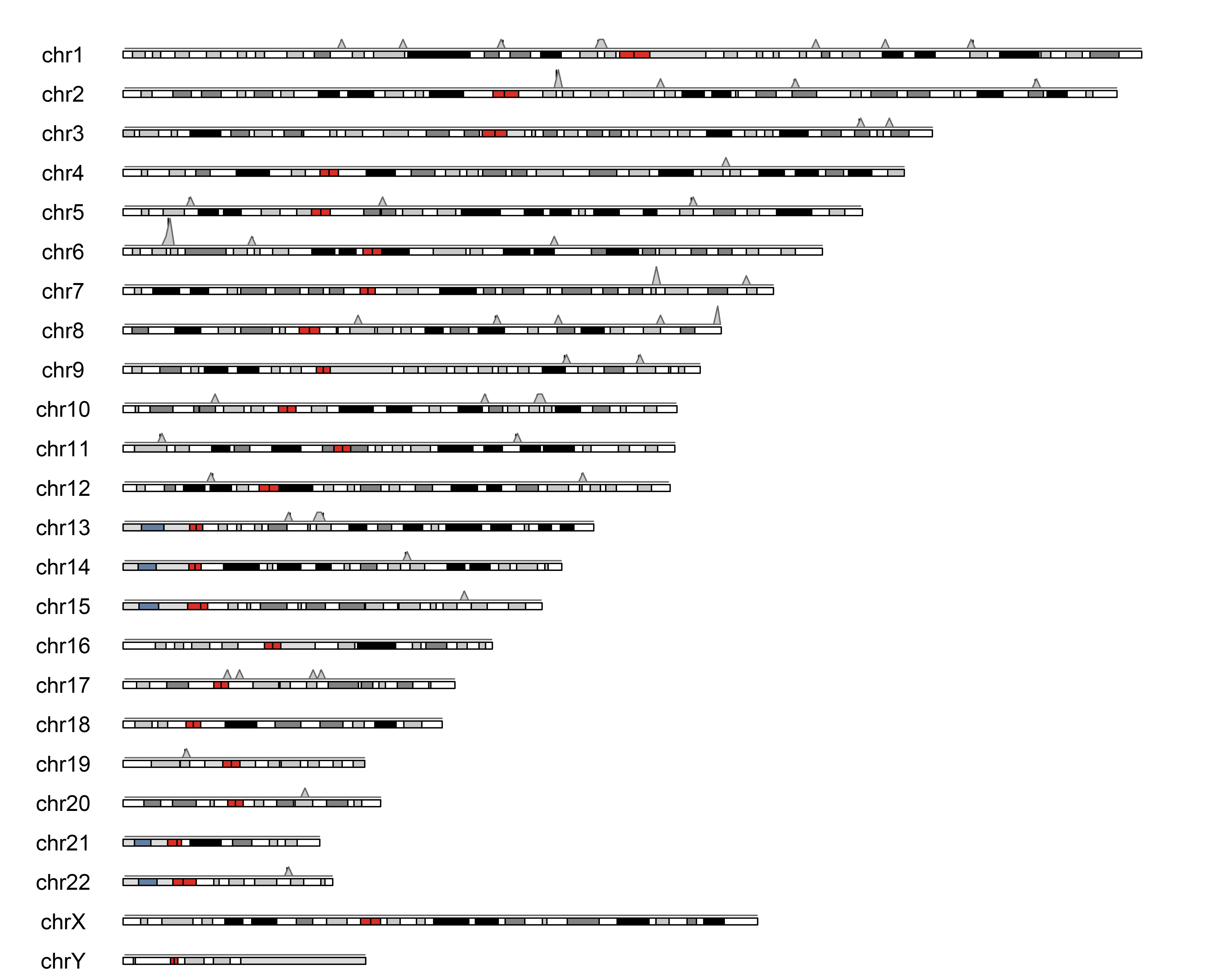

### Supplementary Figure 12: MEAT

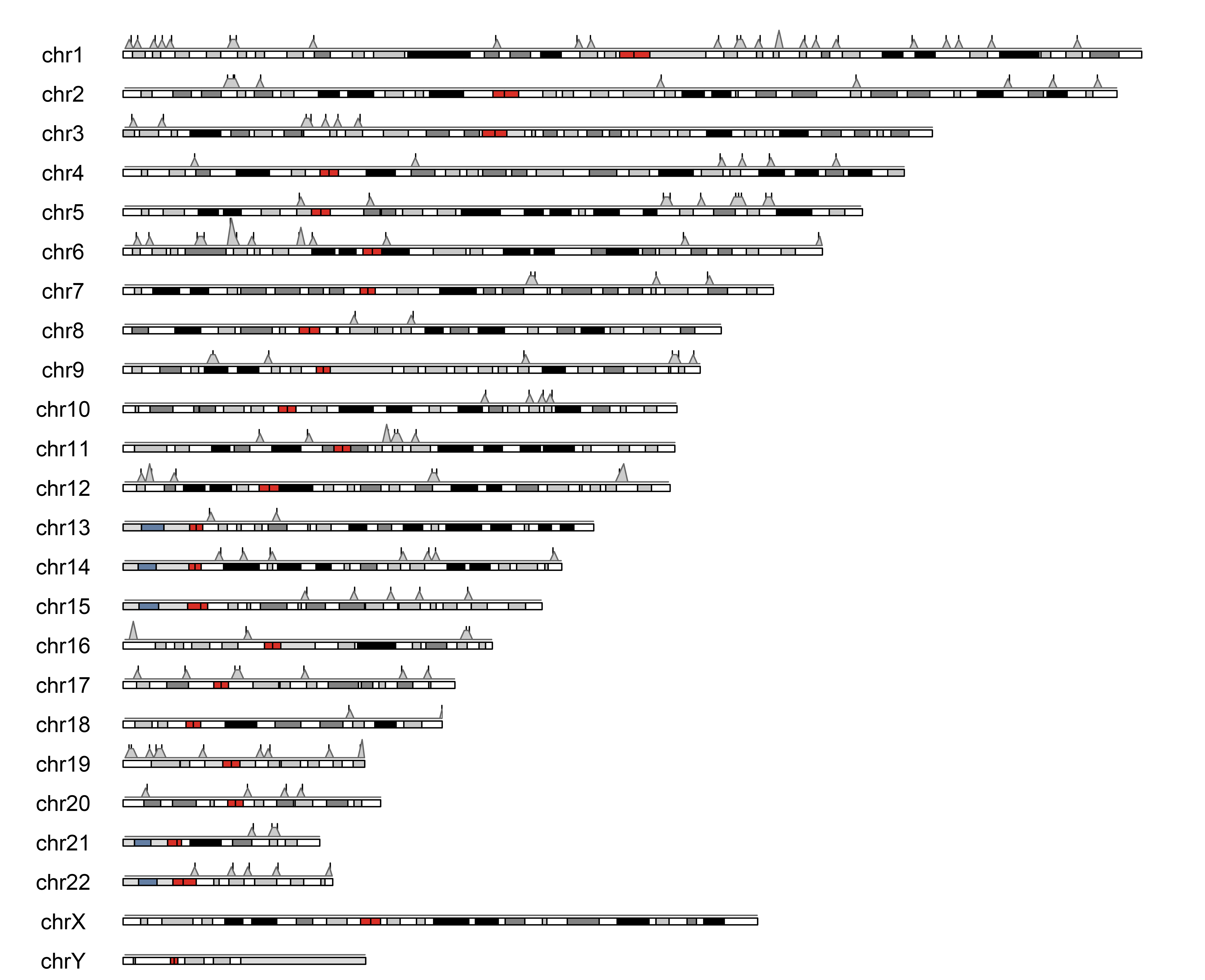

### Supplementary Figure 13: AltumAge

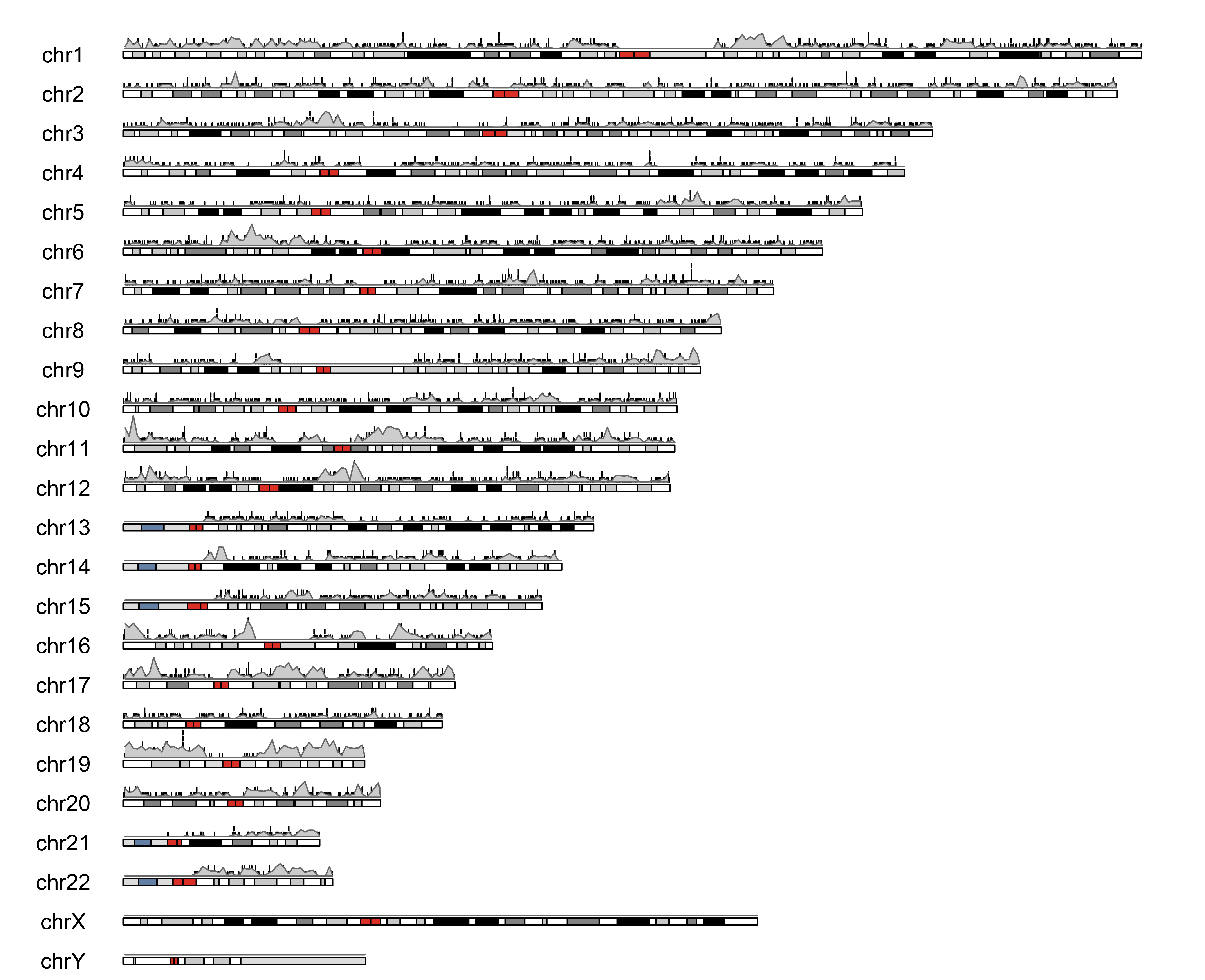

### Supplementary Figure 14: PhenoAge

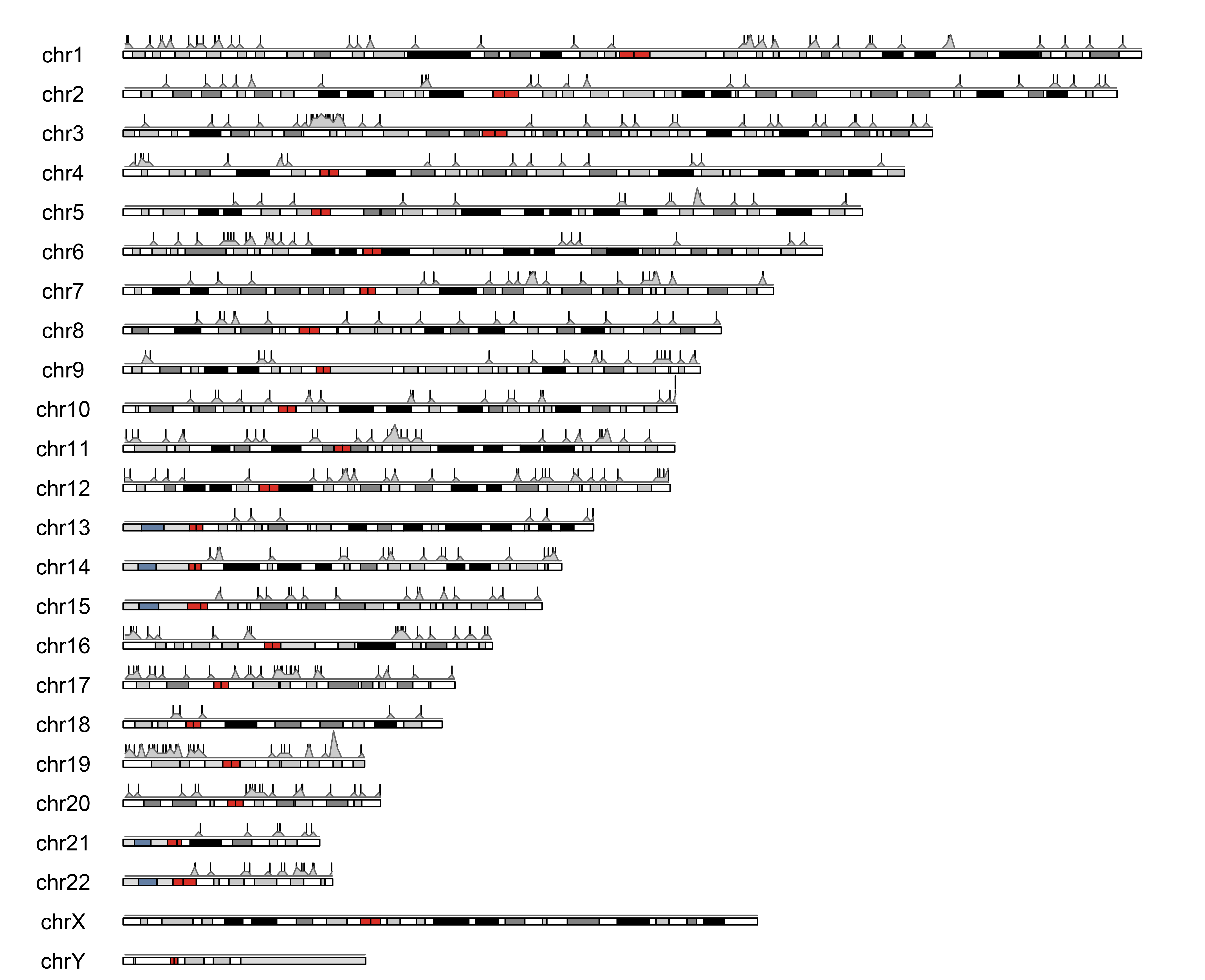

### Supplementary Figure 15: BNN

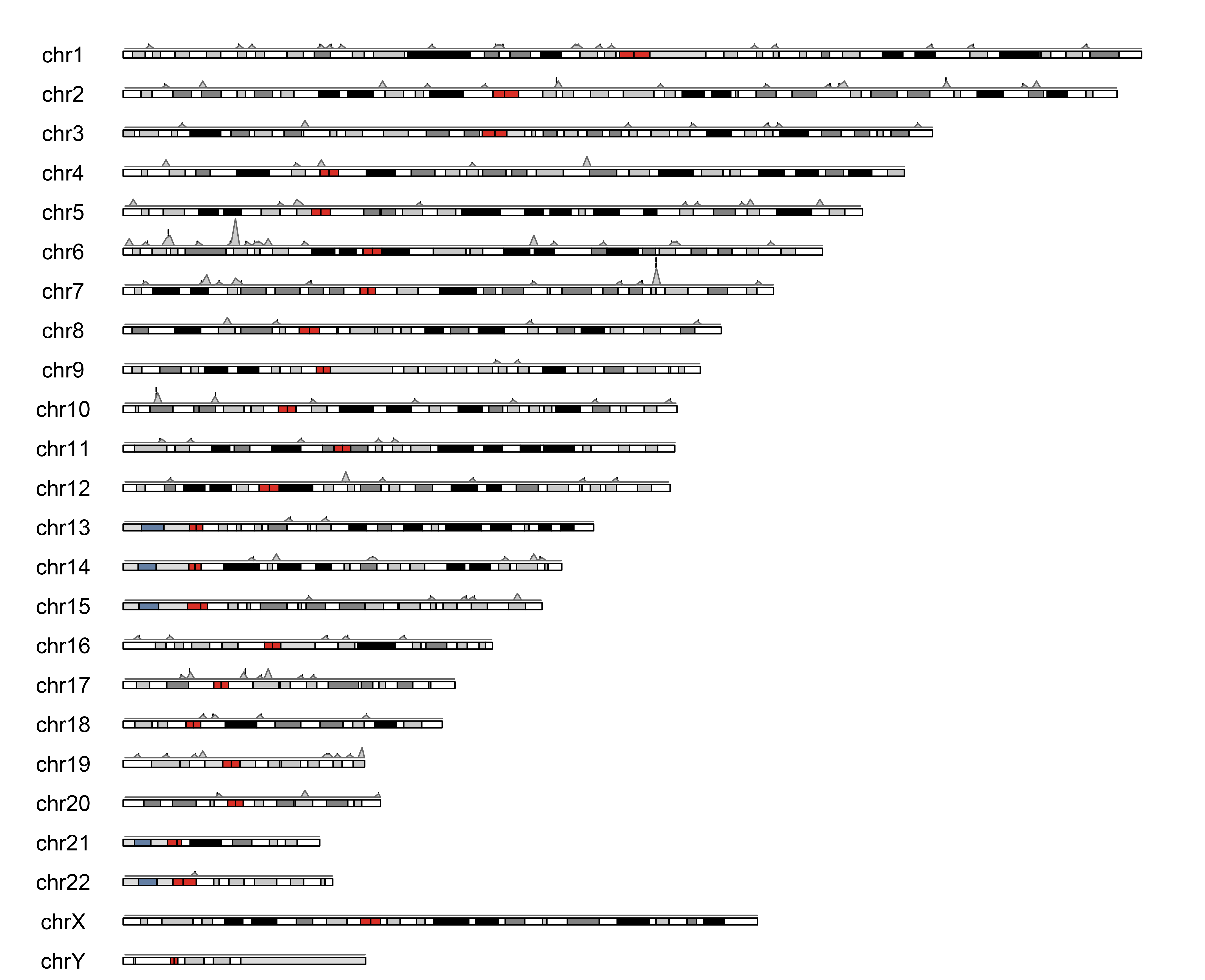

### Supplementary Figure 16: EPM

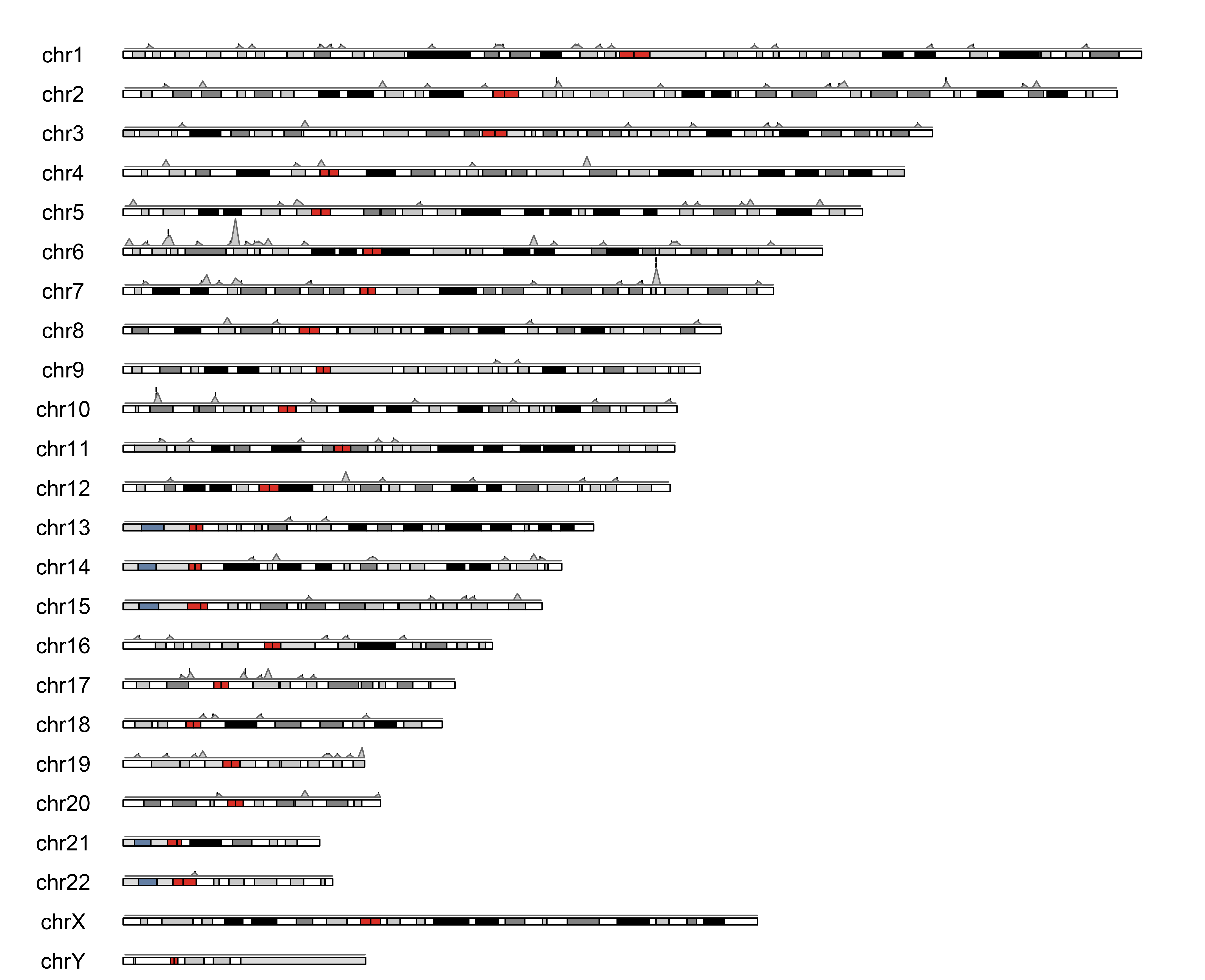

### Supplementary Figure 17: CorticalClock

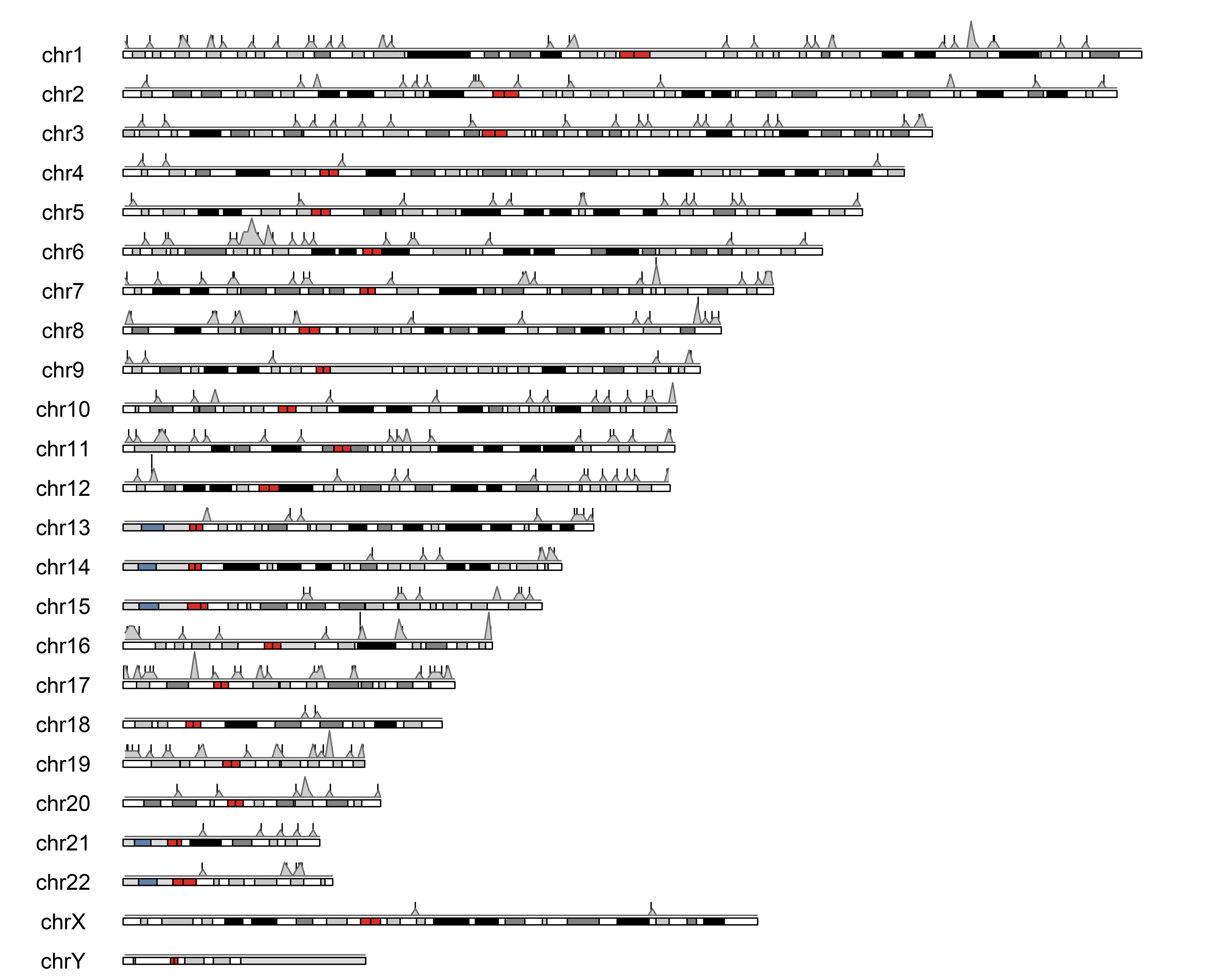

### Supplementary Figure 18: VidalBraloAge

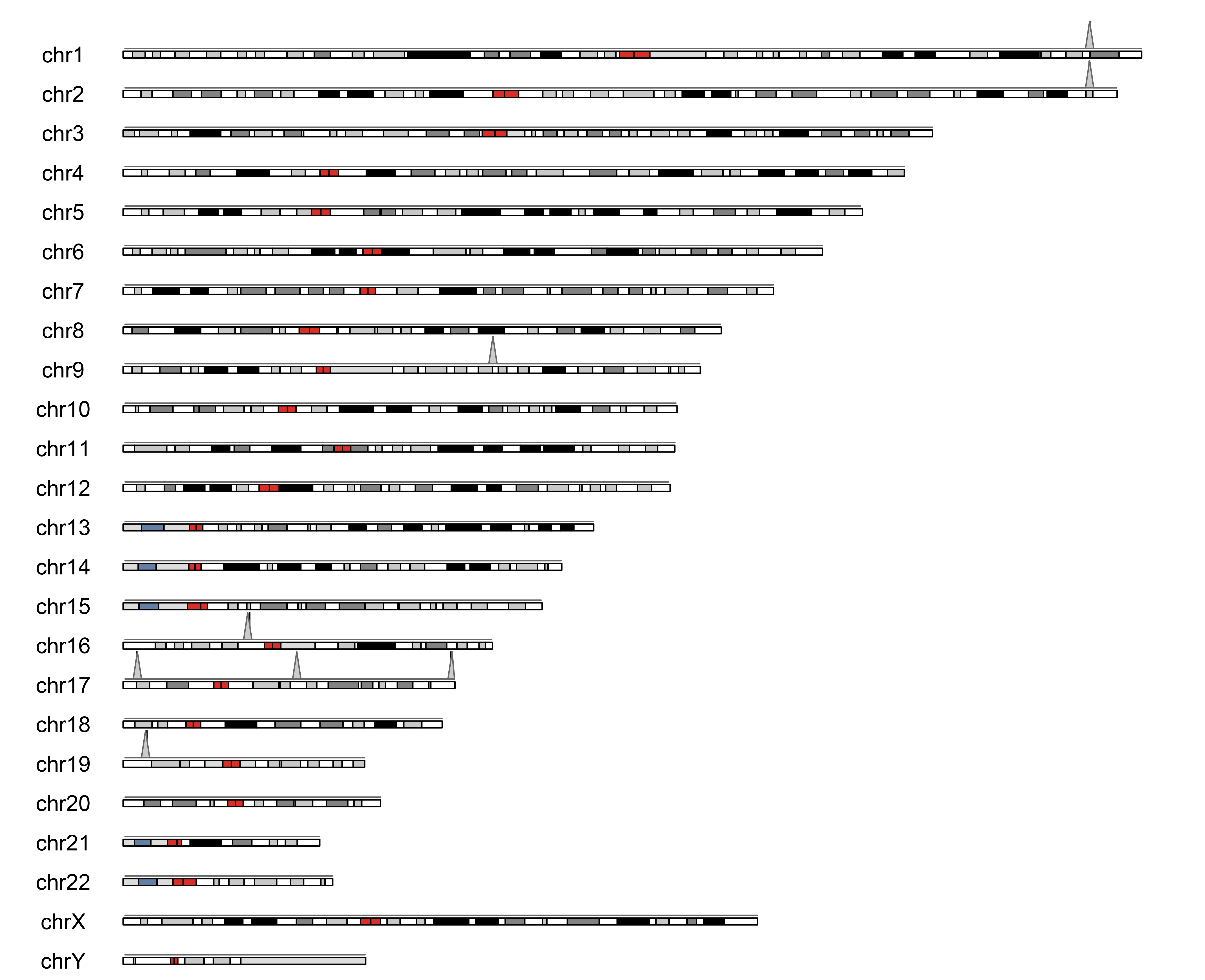

### Supplementary Figure 19: Disease & Healthy

A:

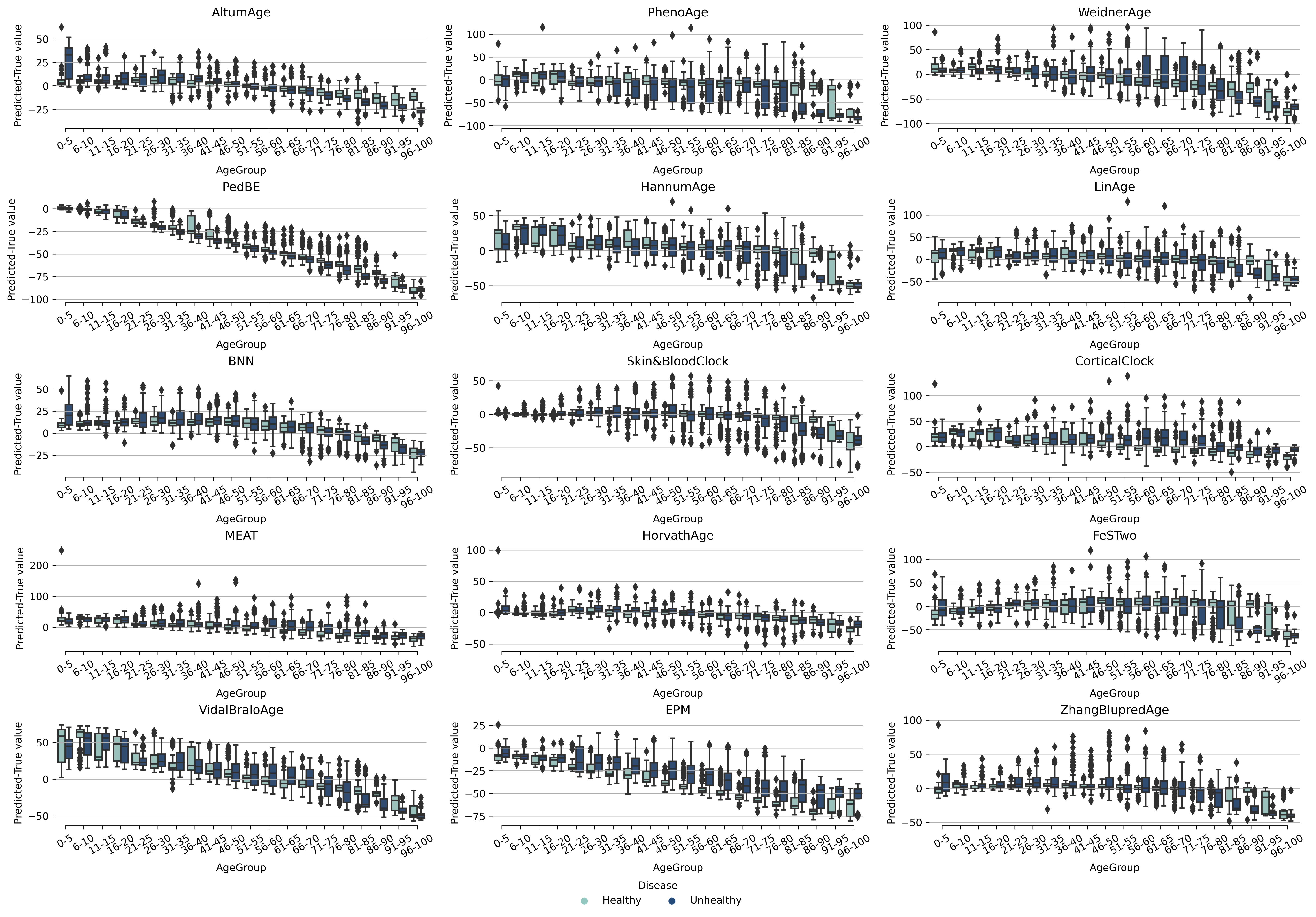

B:

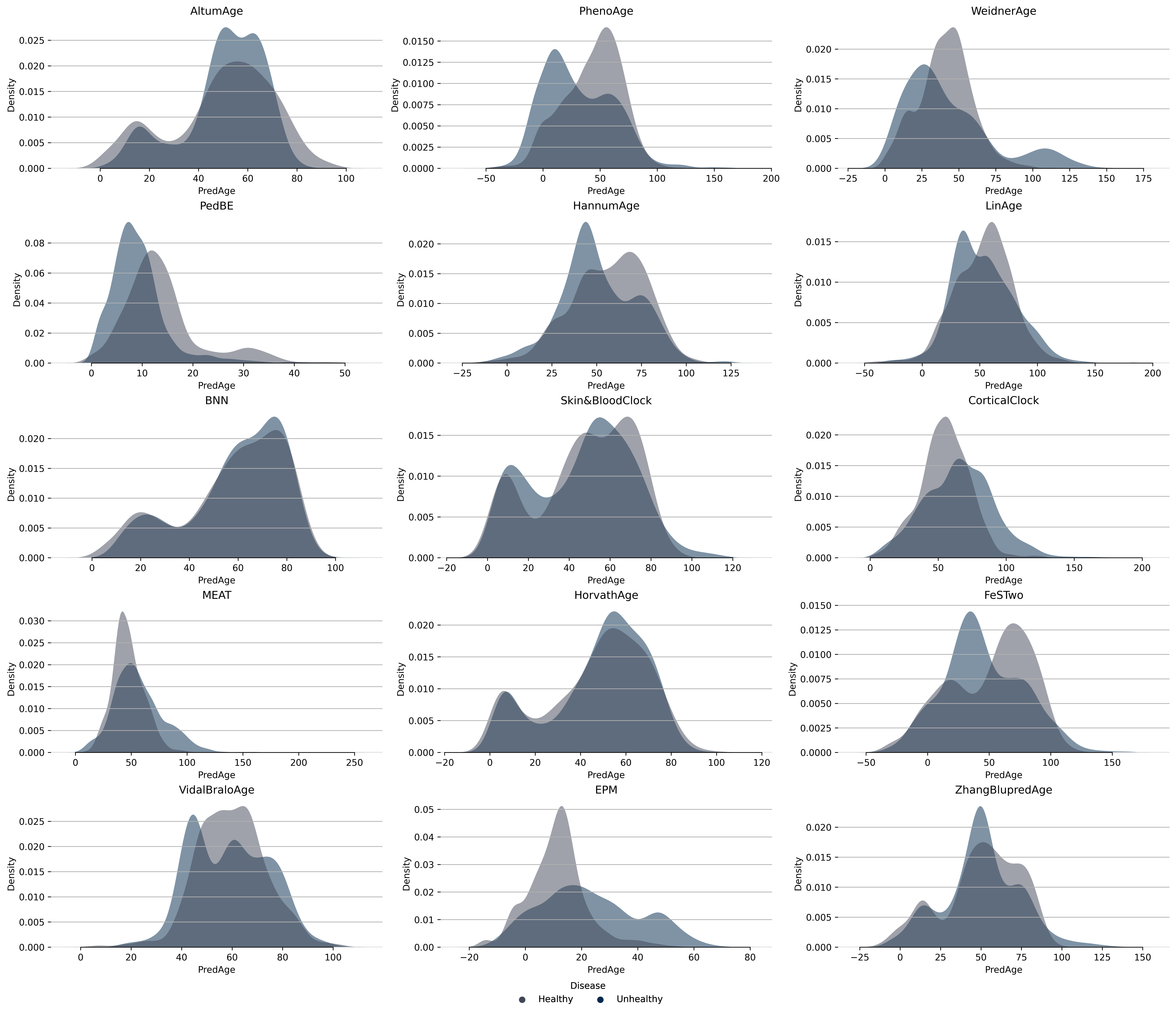

### Supplementary Figure 20: Disease- Parkinsons disease

A:

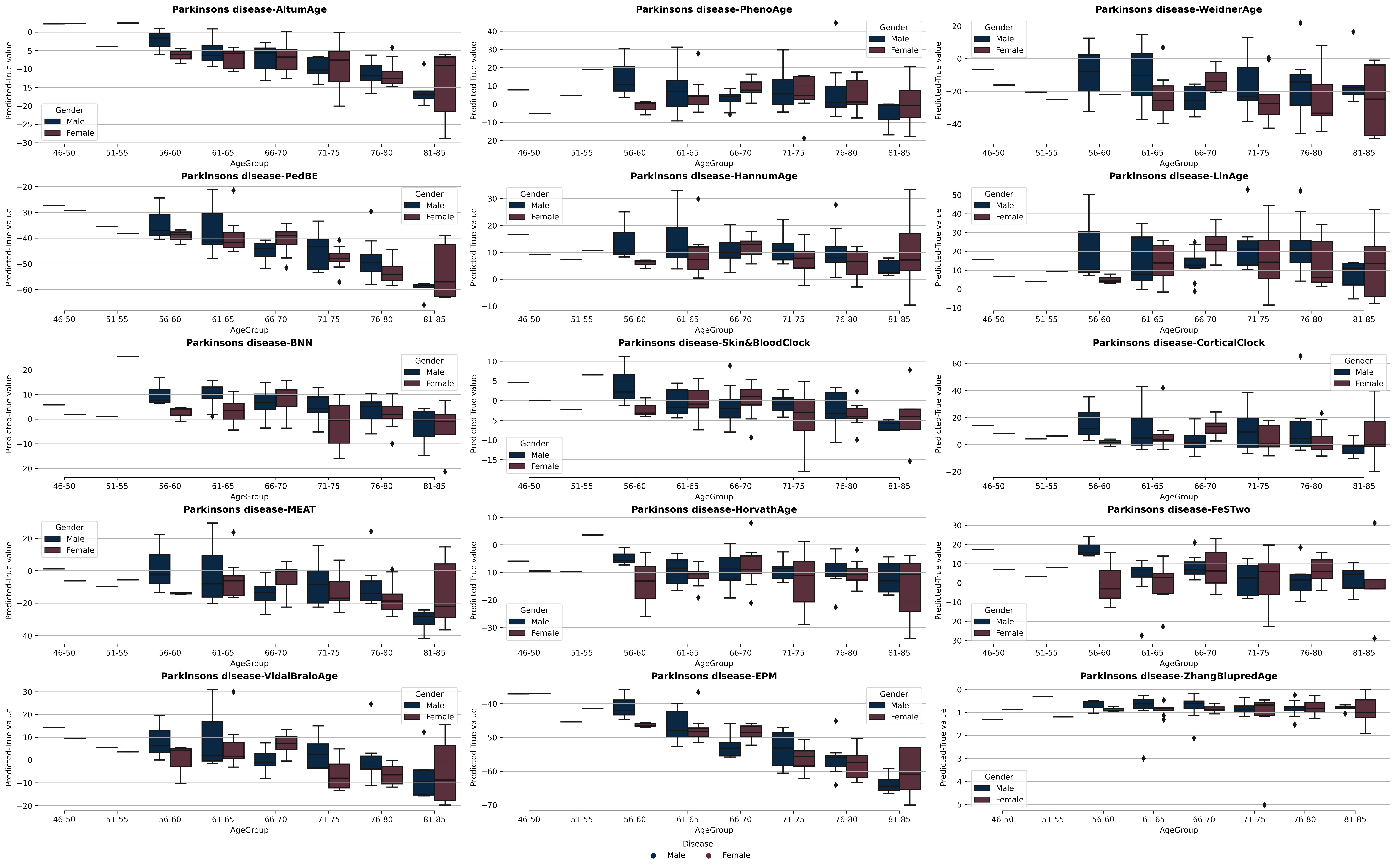

B:

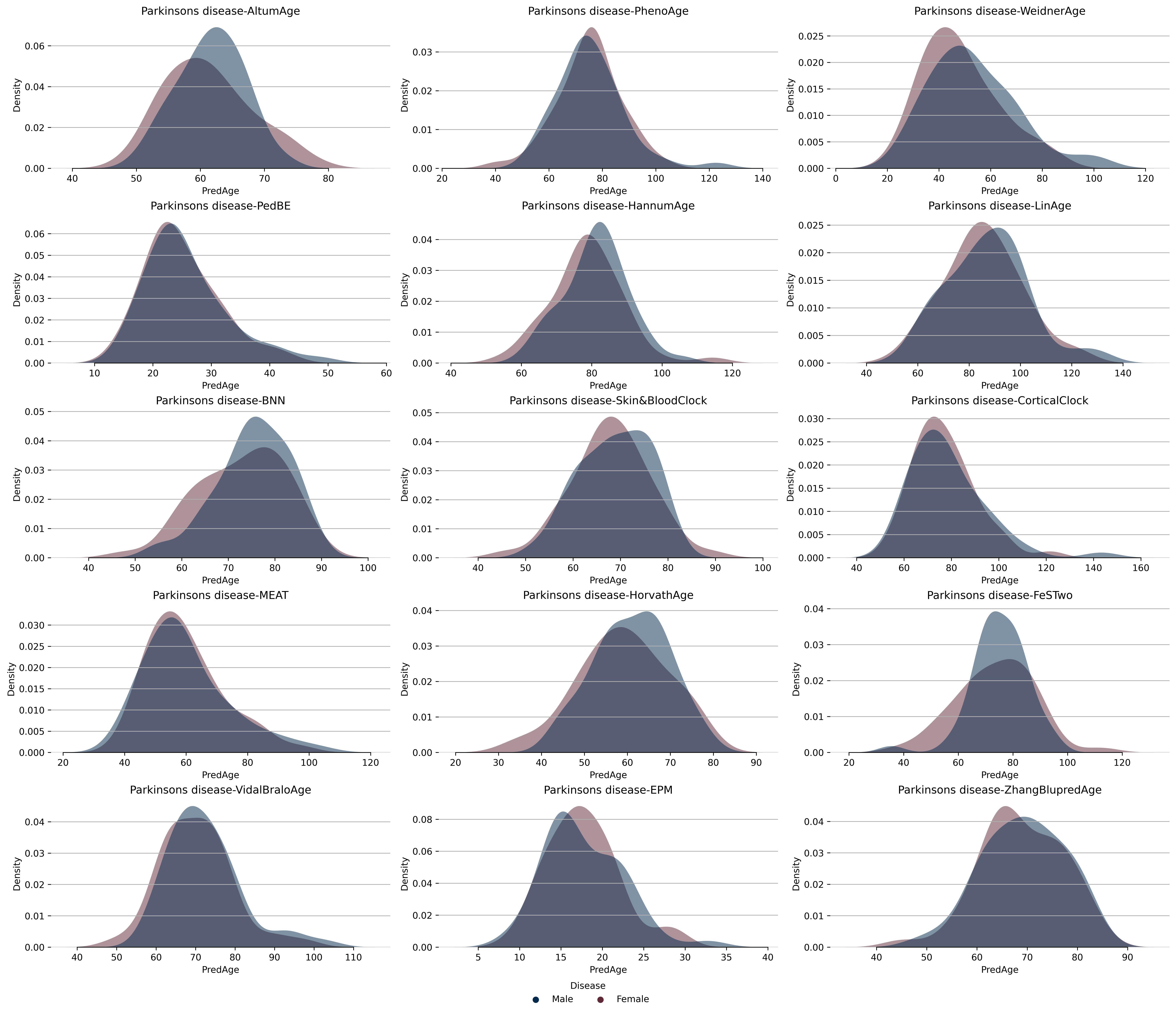

### Supplementary Figure 21: Disease- Small intestine cancer

A:

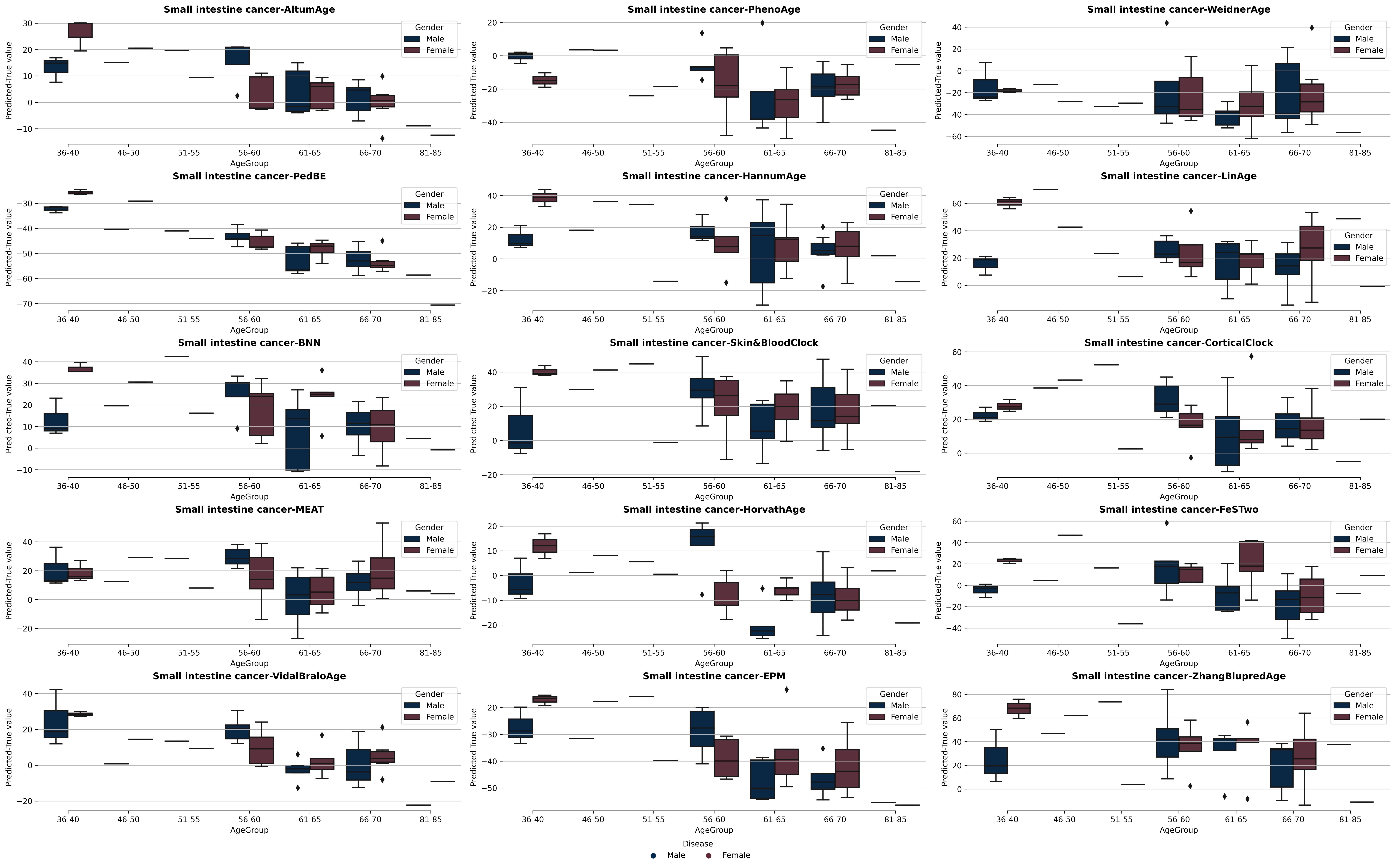

B:

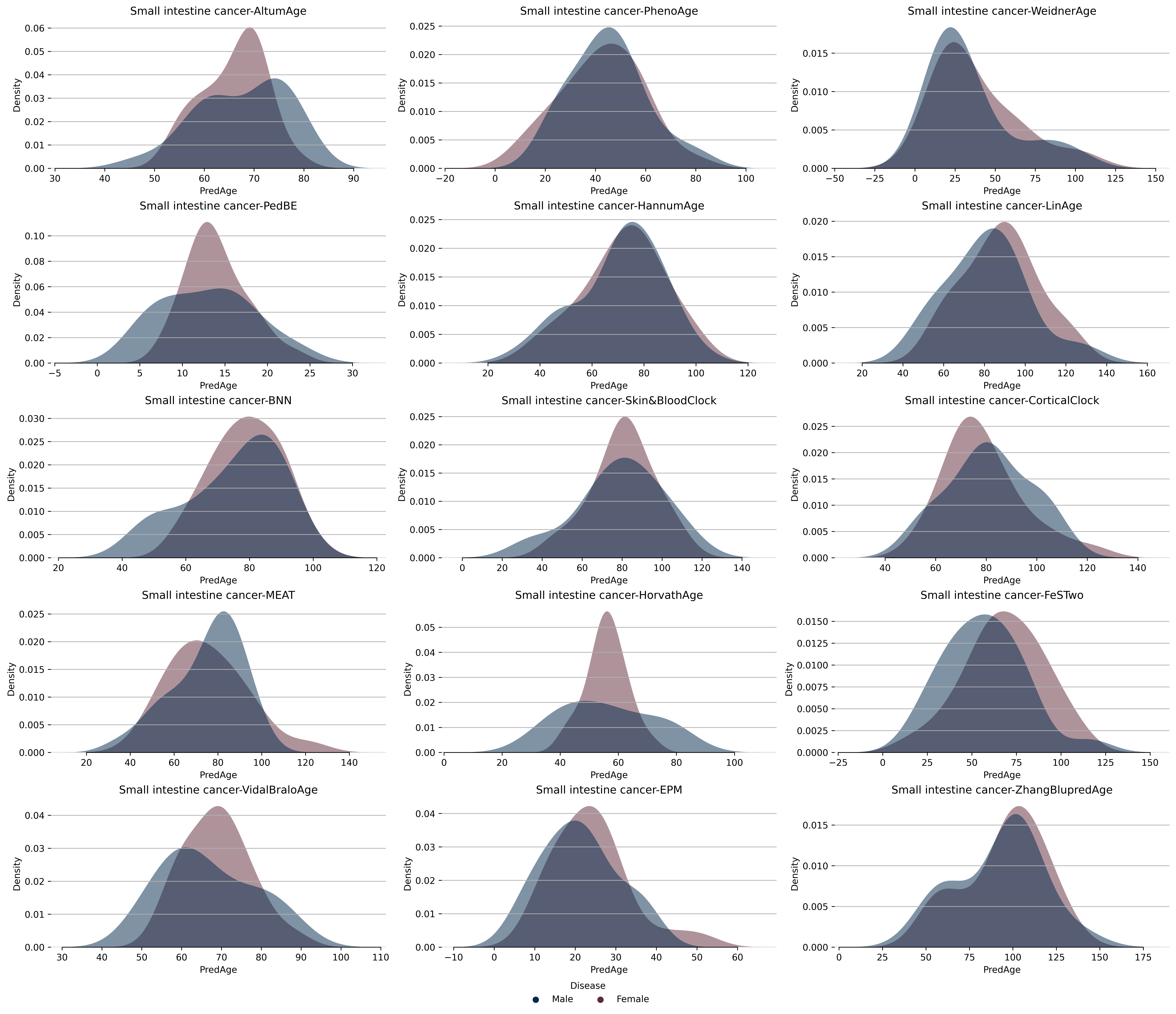

### Supplementary Figure 22: Disease- Colon cancer

A:

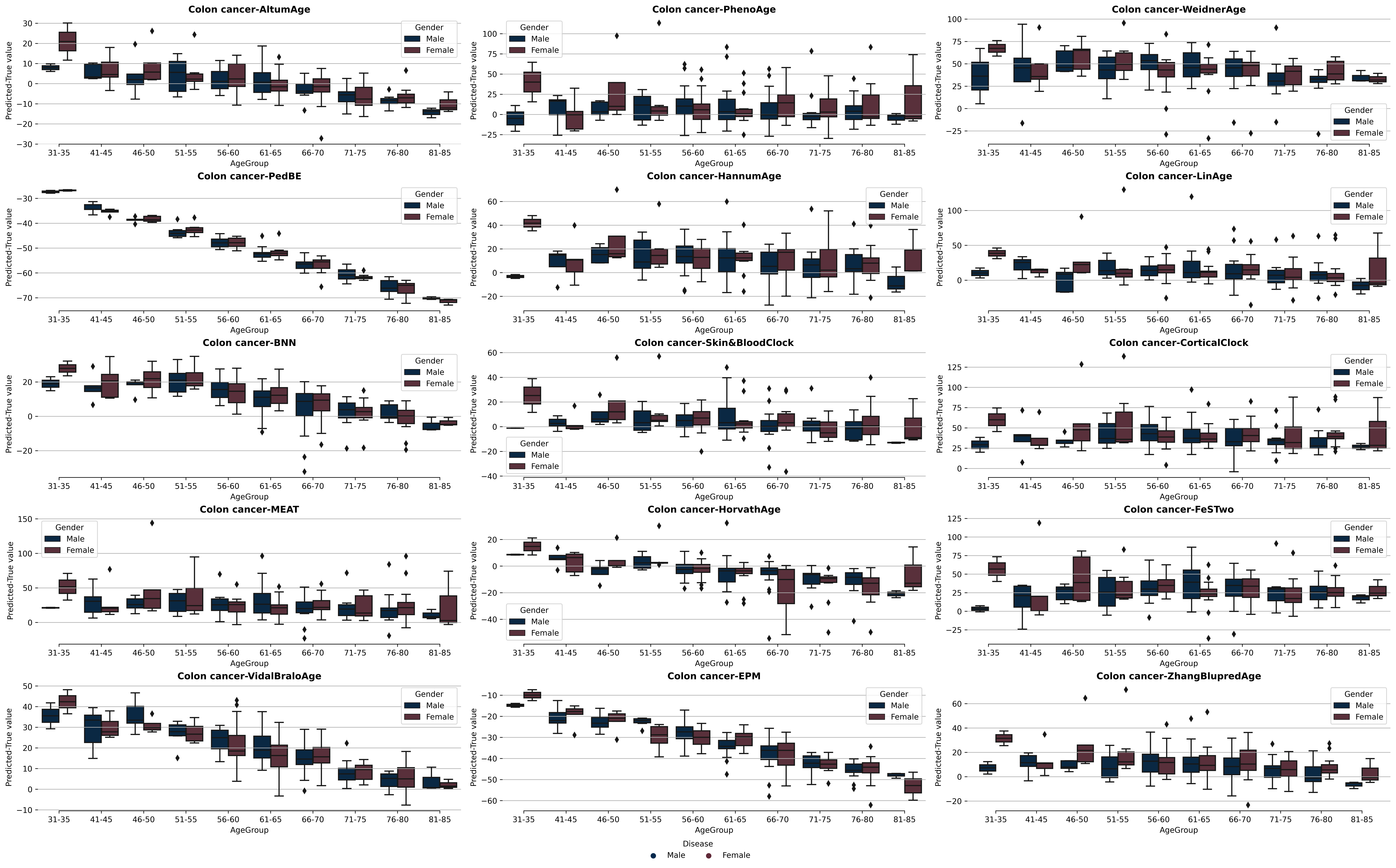

B:

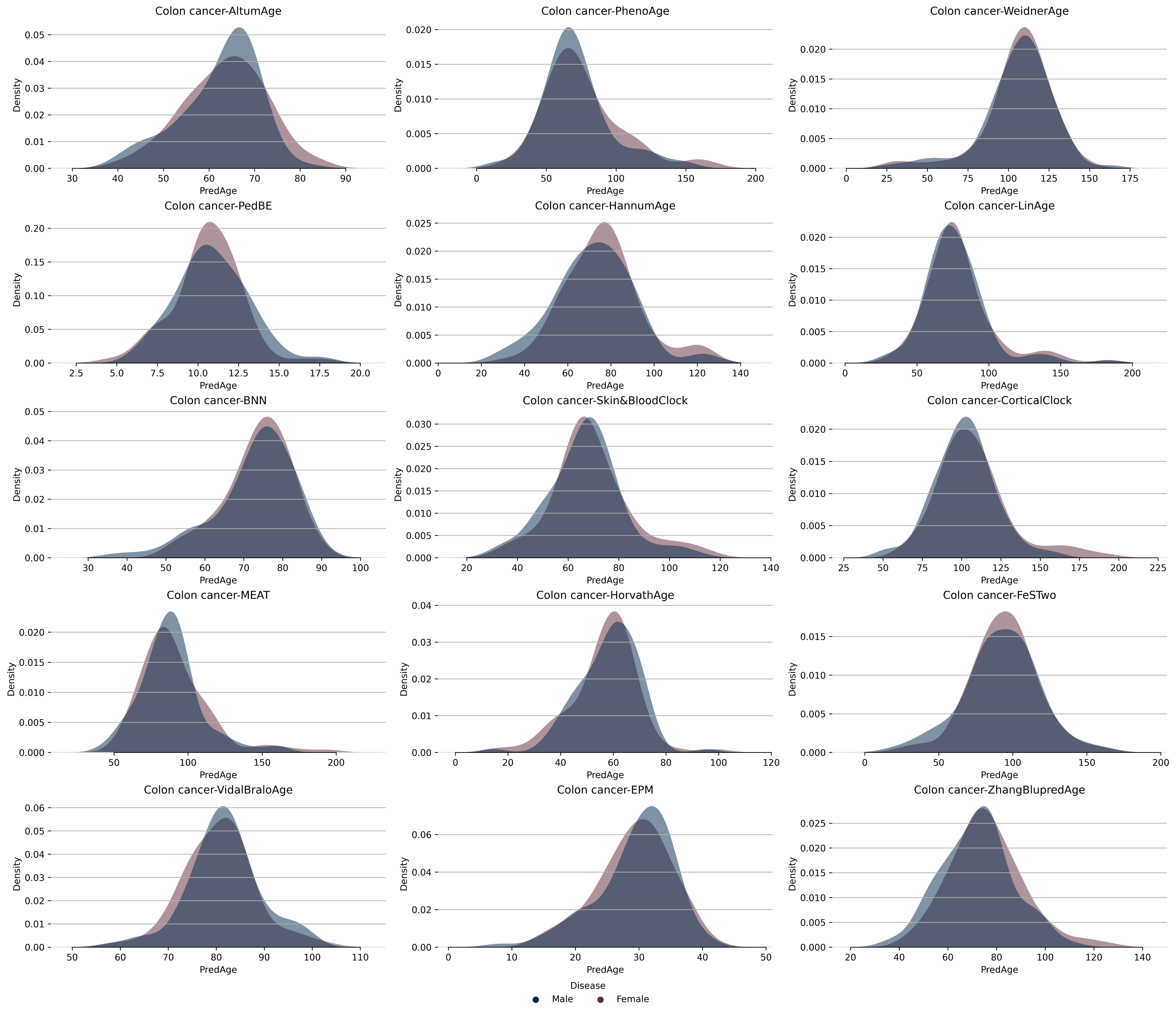

### Supplementary Figure 23: Disease- Fetal alcohol spectrum disorder

A:

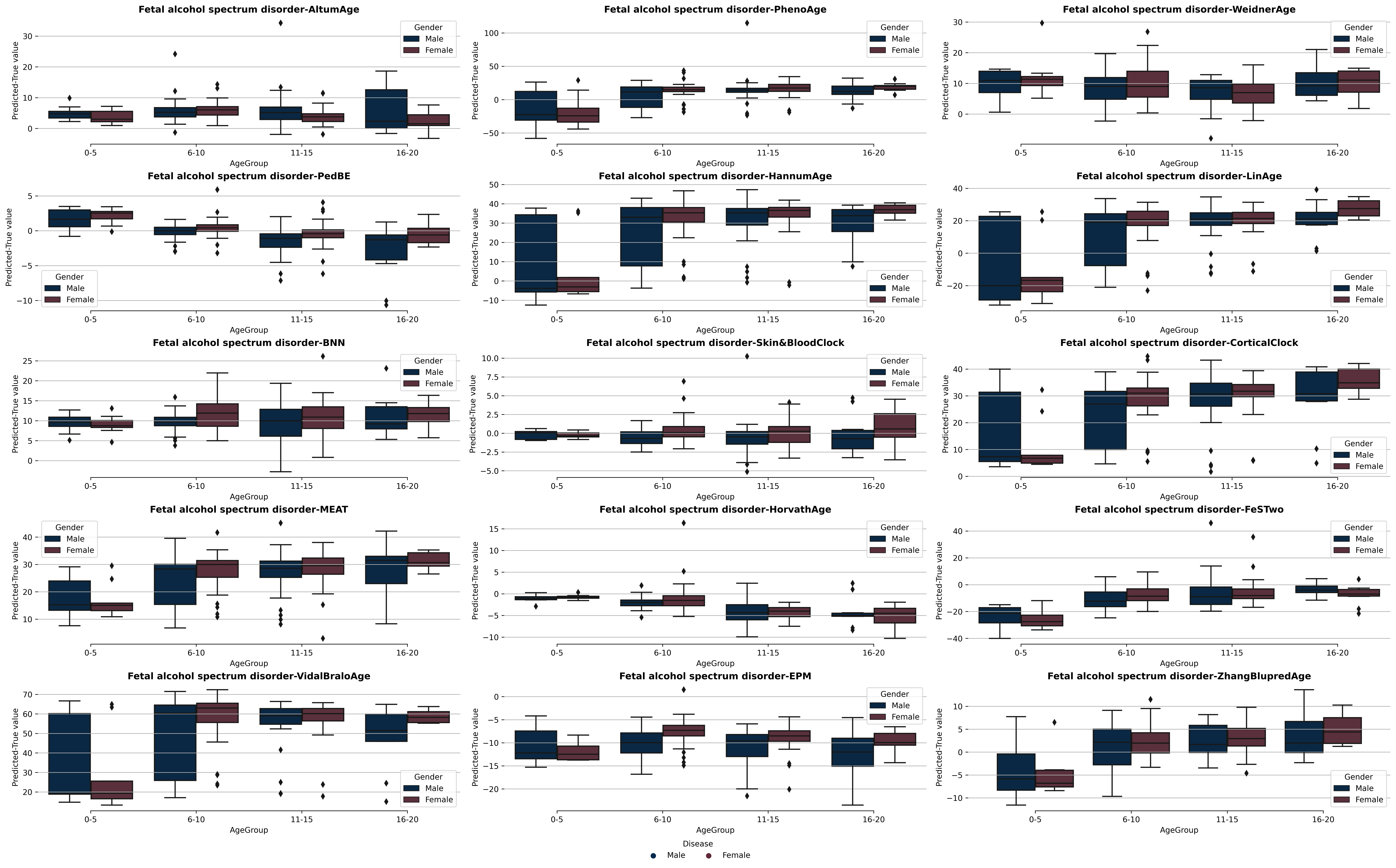

B:

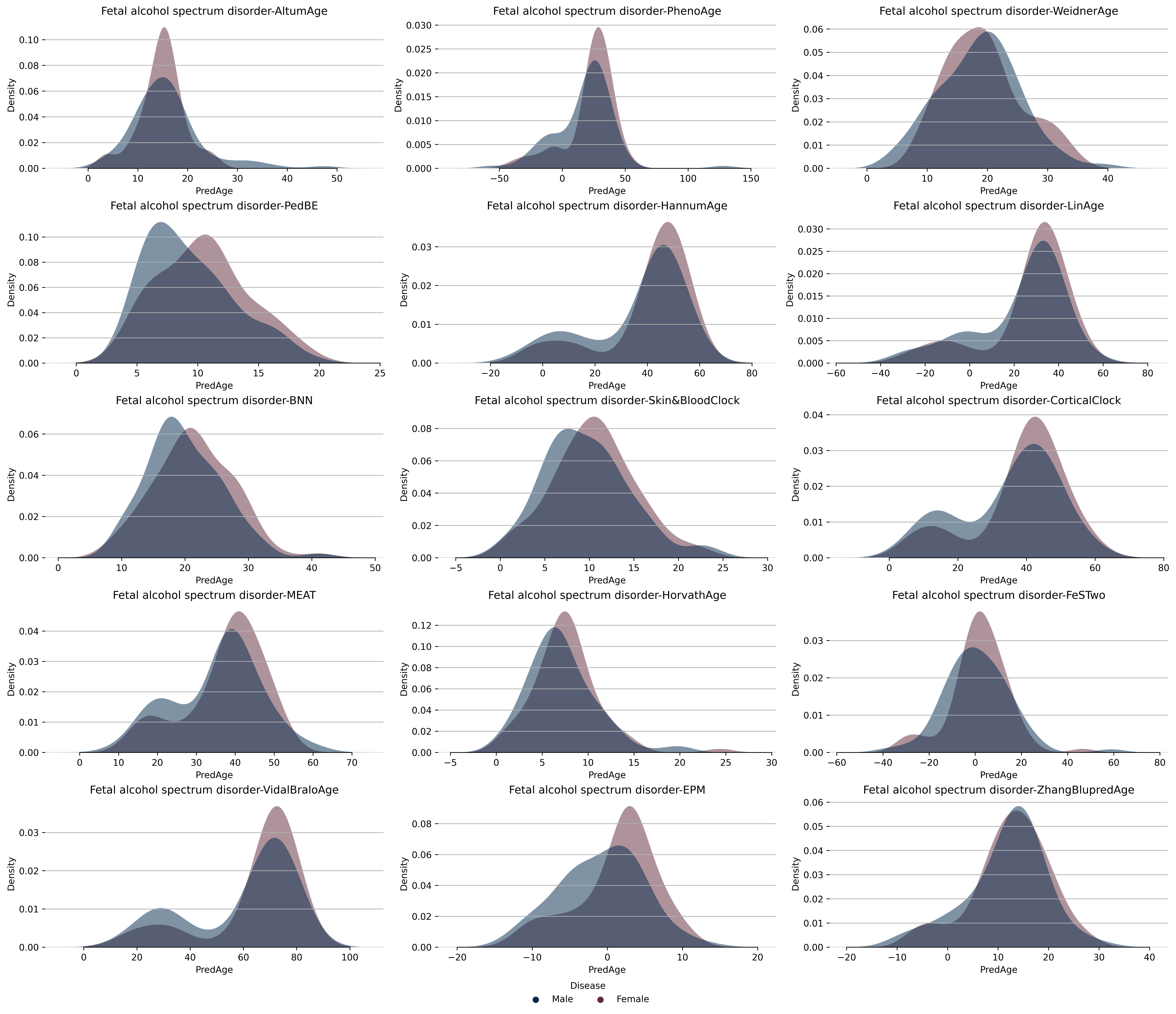

### Supplementary Figure 24: Disease- Alzheimers disease

A:

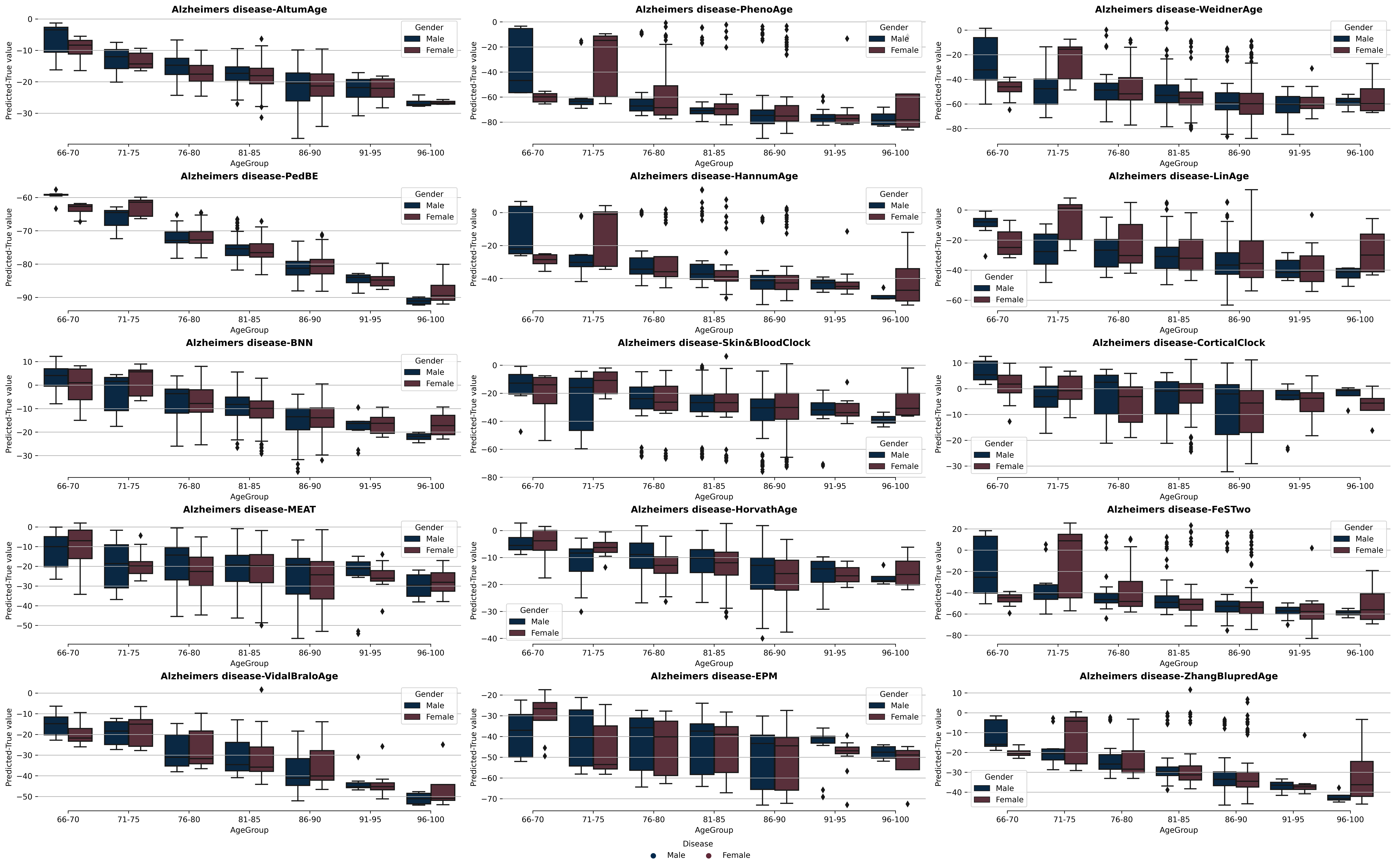

B:

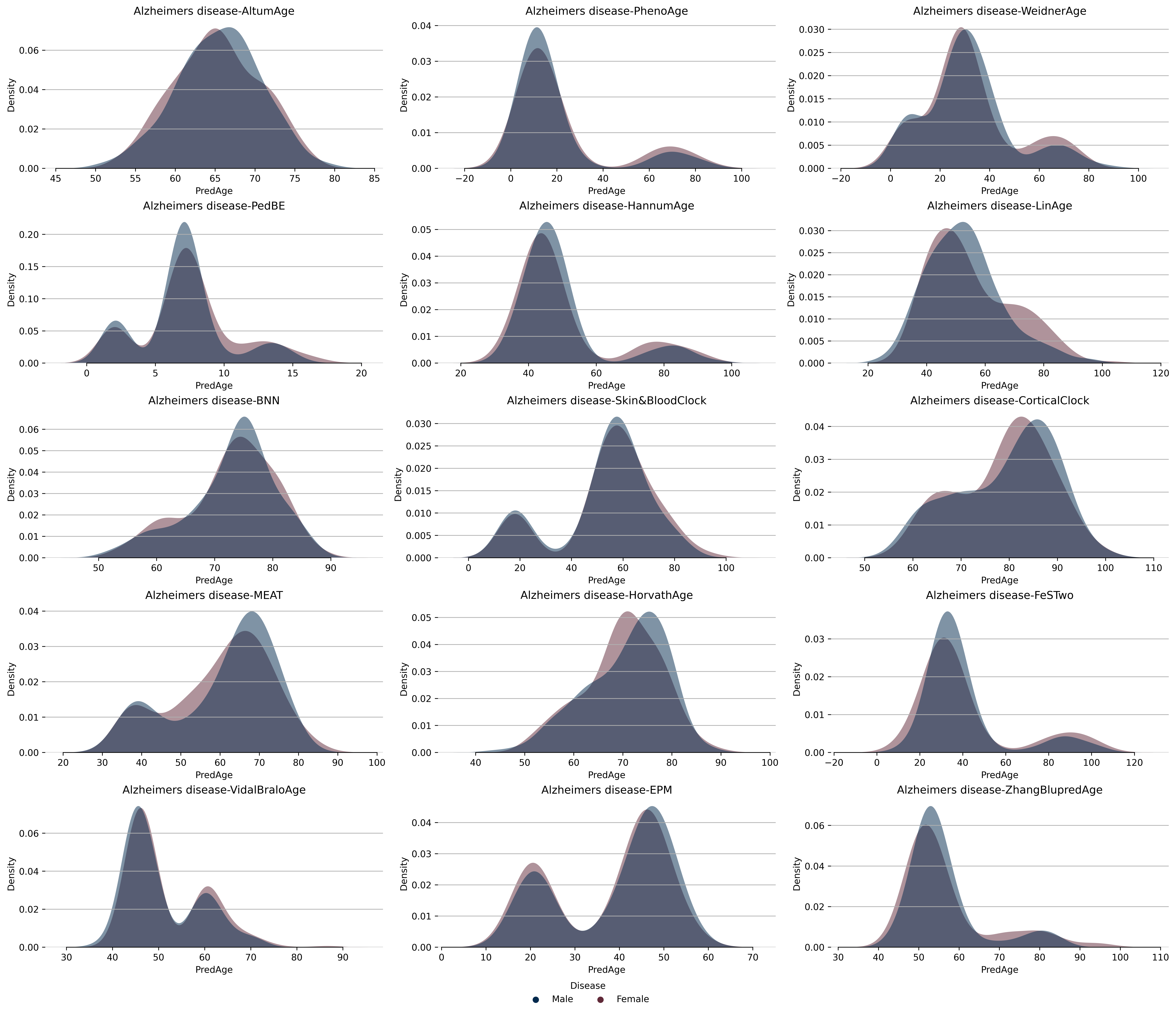

### Supplementary Figure 25 Disease- Schizophrenia

A:

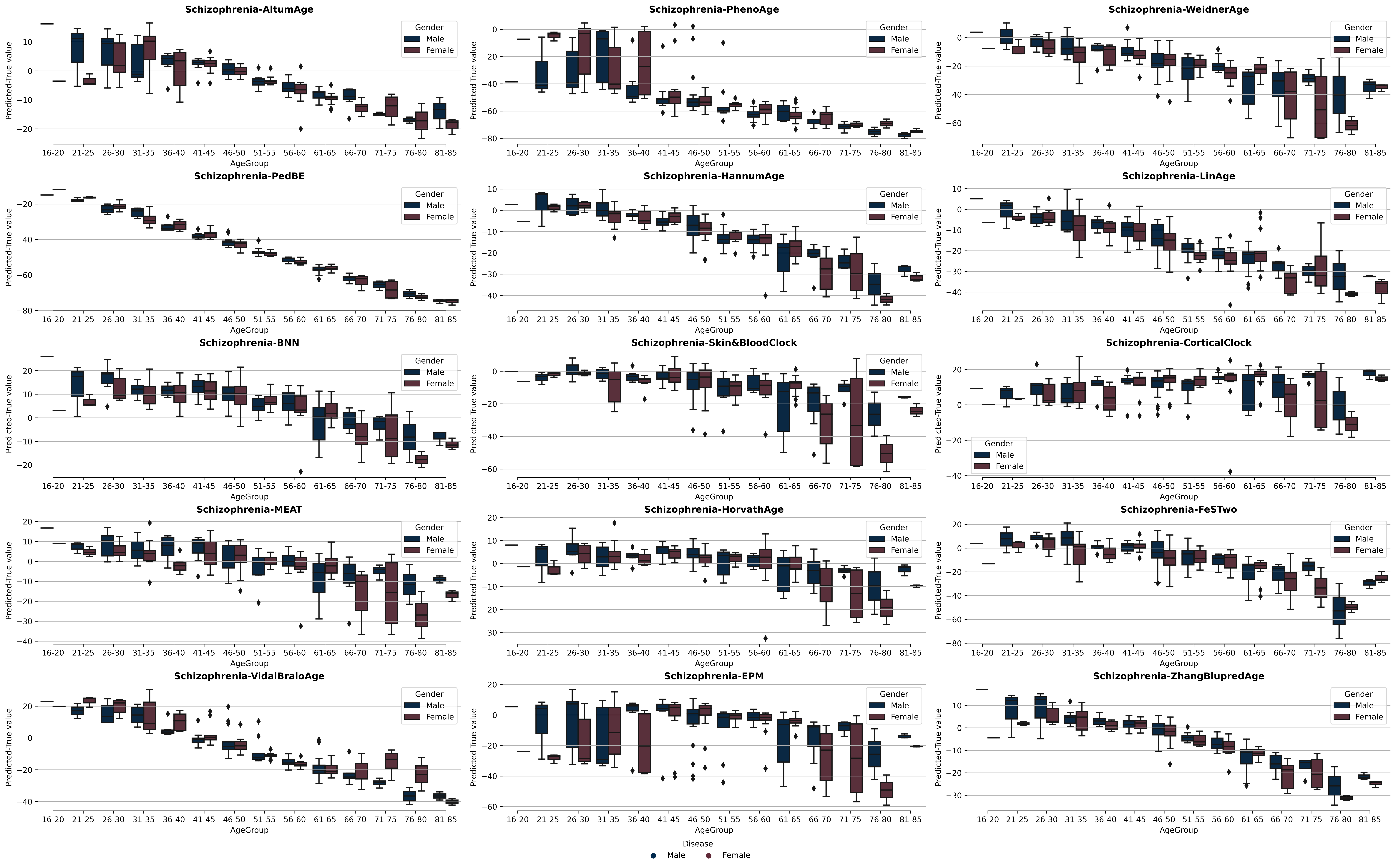

B:

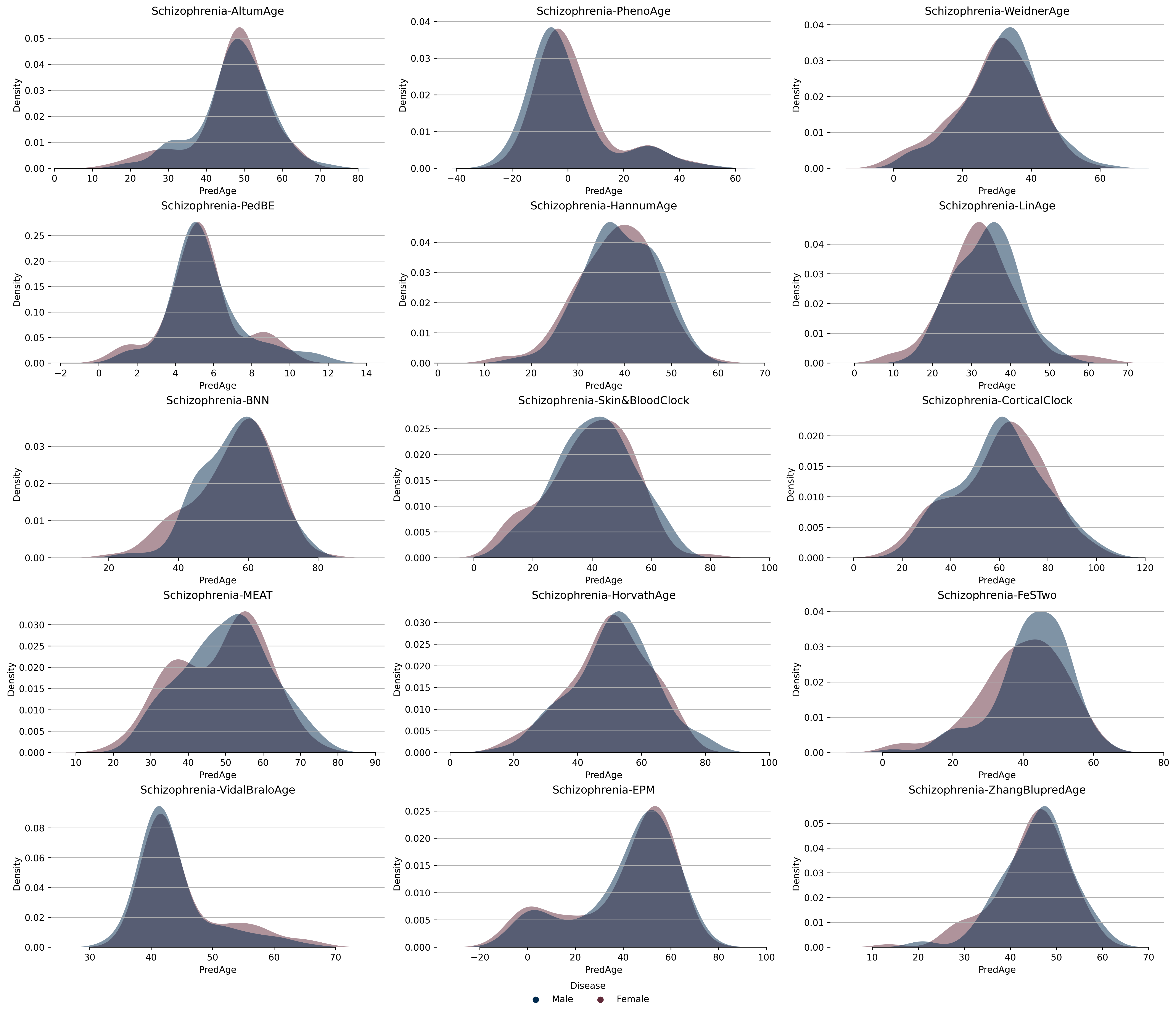

### Supplementary Figure 26 Disease- GSE80970

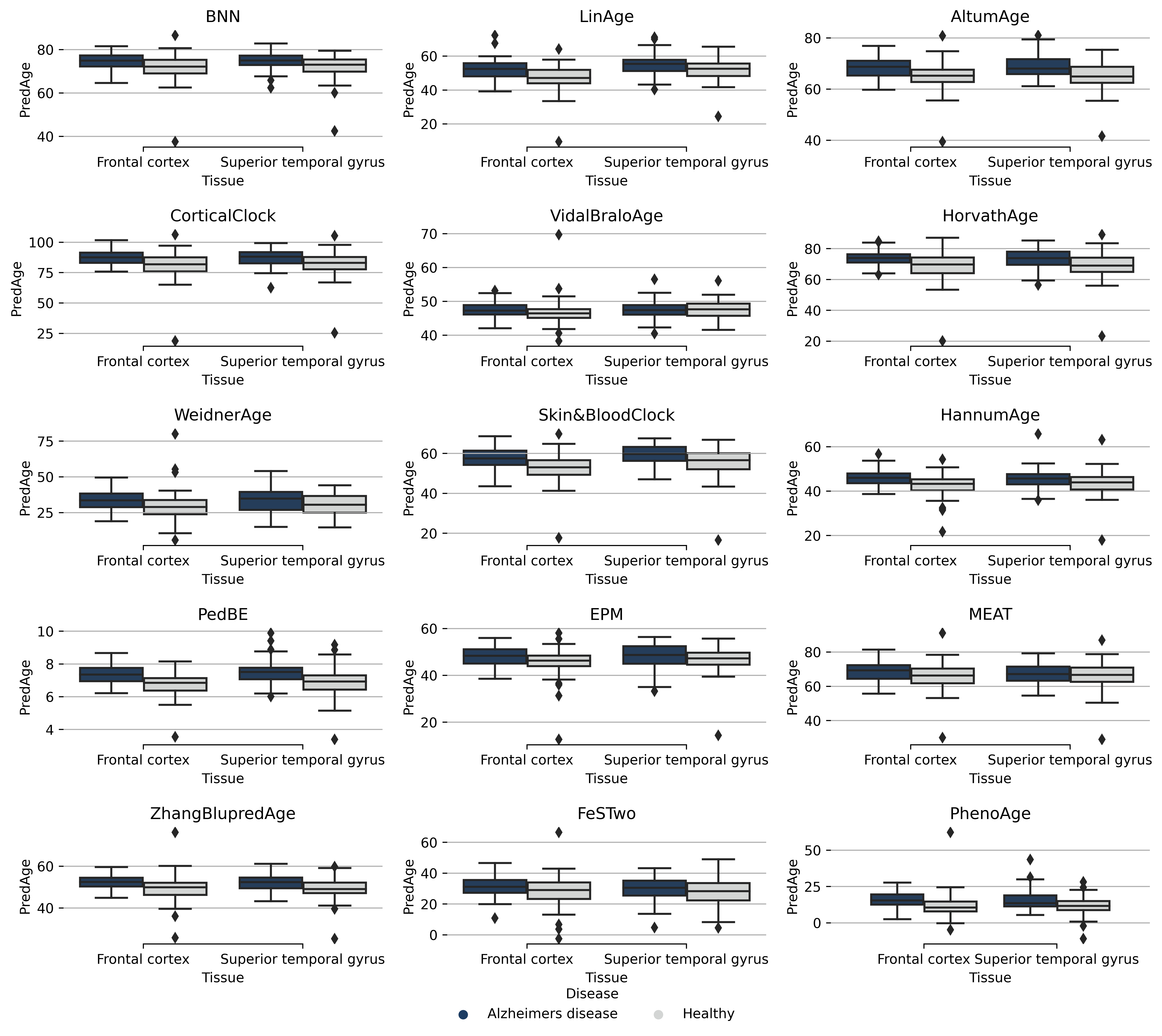

### Supplementary Figure 27 Disease- GSE59685

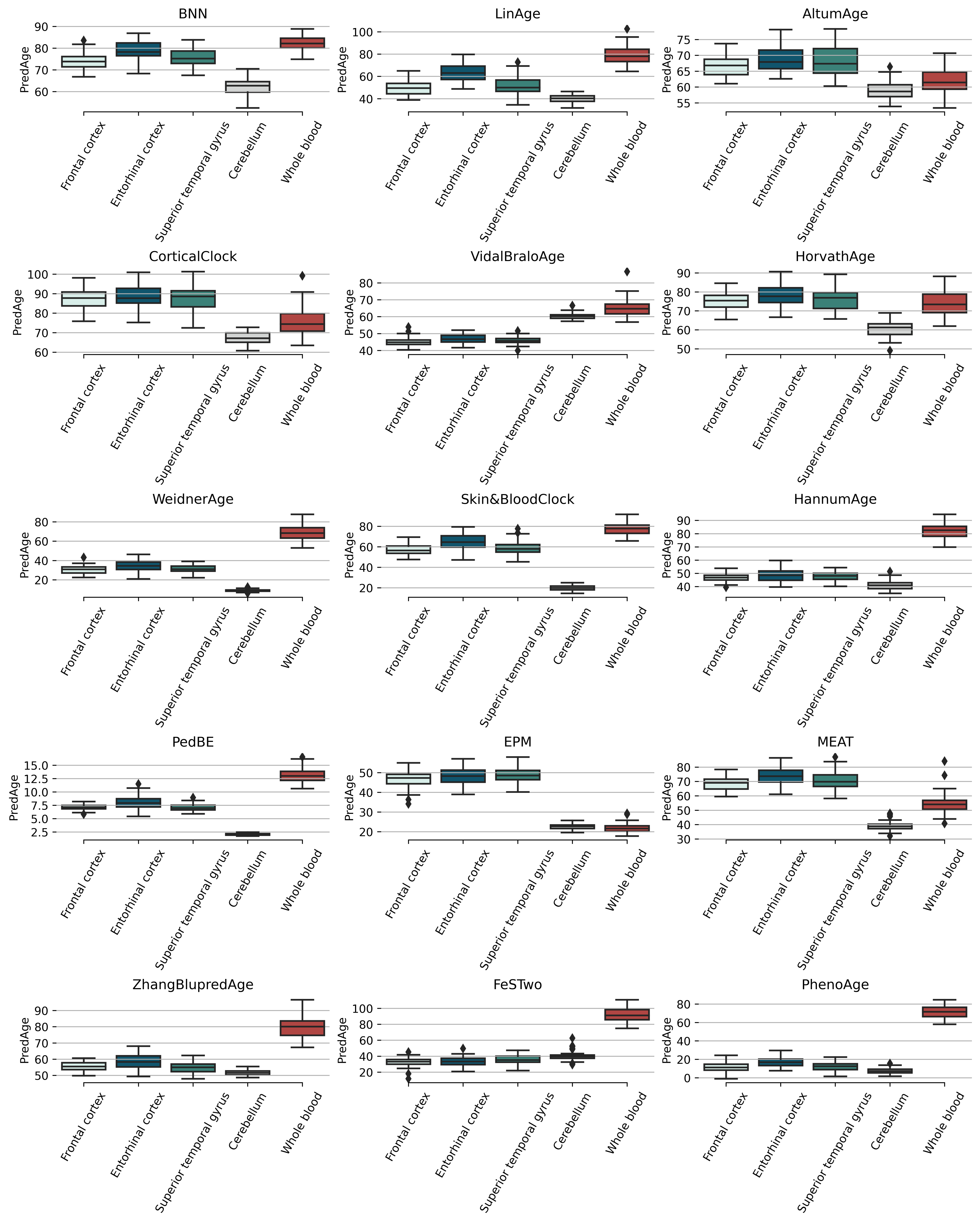
