## Supplementary Note for "A Comprehensive Assessment of Methylation-Based Age Prediction Methods"

### Formula of Expression Matrix Beta

Expression Matrix Beta is used to measure the level of DNA methylation (DNAm). Beta value is actually the percentage of methylation signal intensity. The denominator in the formula is to eliminate the influence of numerical scale and unit, and to avoid the occurrence of zero in the denominator. Its formula is:

$$Beta=\frac{Methylated}{\left( Methylated+Unmethylated+100 \right)}$$

Here, the Methylated represents the signal intensity of methylation, Unmethylated represents the signal intensity of non-methylation.

### Formula of Pearson Correlation Coefficient (PCC)

The PCC is the Pearson correlation coefficient between DNAm age (predicted age) and chronological age (CA). The value of PCC is between in [-1, 1] can measure linear relationship between two sets, which the larger the absolute value has stronger correlation, PCC=-1 has negative correlation, PCC=1 has positive correlation, and PCC=0 does not have linear relationship. It has the following limitations: it cannot reflect the effect of DNAm age calibration, it cannot be calculated when subjects’ CA are same in dataset (that was expressed in NA), and it strongly depends on the standard deviation of age

The Pearson Correlation Coefficient is a method of measuring the linear correlation between two variables. Its formula is:

$$\rho_{X,Y}=\frac{cov(X,Y)}{\sigma_{X}\sigma_{Y}}=\frac{E[\left( X-\mu_{X} \right)\left( Y-\mu_{Y} \right)]}{\sigma_{X}\sigma_{Y}}$$

Here, $\rho_{X,Y}$ represents the correlation coefficient between X and Y; $cov(X,Y)$ represents the covariance between X and Y, and $\sigma_{X}, \sigma_{Y}$ represent the standard deviation of X and Y, respectively.

### Formula of Median Absolute Error (MAE)

The MAE is the median absolute difference between sets of DNAm age and CA. It can measure the stability of model prediction and the error is well suited for studying whether DNAm age is poorly calibrated. The value of MAE is greater than 0. The closer the value is to 0, the closer the DNAm age and CA are. It is sensitive to outliers.

The Median Absolute Error (MAE) is a metric used to evaluate the performance of regression models. It is defined as the median of the absolute differences between the predicted and actual values. The formula for MAE is:

$$MAE=\frac{1}{n}\sum_{i=1}^{n} |y_{i}-\hat{y}_{i}|$$

Here, $y_{i}$ represents the actual value of the $i^{th}$ observation, $\hat{y}_{i}$ represents the predicted value of the $i^{th}$ observation, and n represents the total number of observations.

### Formula of Root Mean Square Error (RMSE)

The RMSE is Root Mean Square Error between sets of DNAm age and CA. The value of RMSE is greater than 0. The closer the value is to 0, the closer the DNAm age and CA are. Compare with MAE, RMSE reduced the sensitivity to unit dimensions and the error is magnified (as described below).

The Root Mean Square Error (RMSE) is a metric used to evaluate the performance of regression models. It measures the square root of the average of the squared differences between the predicted and actual values. The formula for RMSE is:

$$RMSE=\sqrt{\frac{1}{n}\sum_{i=1}^{n} {(y_{i}-\hat{y}_{i})}^{2}}$$

Here, $y_{i}$ represents the actual value of the $i^{th}$ observation, $\hat{y}_{i}$ represents the predicted value of the $i^{th}$ observation, and n represents the total number of observations.

### Score aggregation

To rank methods, we aggregated the ranked scores across five criteria. The RMSE score was selected to evaluate the powerful of accuracy, generality, and the effect of missing values. The time and memory metrics were applied in the scalability. We self-define the usability scores by weight. Each criterion contains many contents that were calculated by individual metric and ranked, the ranks were aggregated as the final criterion score. The overall score aggregated the five criteria’s rank.
